# *In situ* differentiation of iridophore crystallotypes underlies zebrafish stripe patterning

**DOI:** 10.1101/2020.03.25.008664

**Authors:** Dvir Gur, Emily Bain, Kory Johnson, Andy J. Aman, Amalia Pasoili, Jessica D. Flynn, Michael C. Allen, Dimitri D. Deheyn, Jennifer C. Lee, Jennifer Lippincott-Schwartz, David Parichy

## Abstract

Skin color patterns are ubiquitous in nature, evolve rapidly, and impact social behavior^1^, predator avoidance^2^, and protection from ultraviolet irradiation^3^. A leading model system for vertebrate skin patterning is the zebrafish^4-7^; its alternating blue stripes and yellow interstripes depend on guanine crystal-containing cells called iridophores that reflect light. It was suggested that the zebrafish’s alternating color pattern arises from a single type of iridophore migrating differentially to stripes and interstripes^7-9^. When we tracked iridophores, however, we found they did not migrate between stripes and interstripes but instead differentiated and proliferated in place based on their micro-environment. RNA seq analysis further revealed stripe and interstripe iridophores had different transcriptomic states, while cryogenic scanning electron microscopy and micro-X-ray diffraction showed they had different guanine crystal organizations and responsiveness to norepinephrine, all indicating that stripe and interstripe iridophores are different cell types. Based on these results, we present a new model of skin patterning in zebrafish in which distinct iridophore crystallotypes containing specialized, physiologically responsive, subcellular organelles arise in stripe and interstripe zones by *in situ* differentiation. In this model, pattern phenotype depends not only on interactions among pigment cells that affect their arrangements, but also on factors that specify subcellular organization and physiological responsiveness of specialized organelles.

## Introduction

Biological patterning is ubiquitous in nature, but mechanisms underlying its establishment and maintenance have been well-documented in only a few instances that are unlikely to represent the full spectrum of pattern-forming systems^6,10^. Indeed, patterning can arise in response to graded positional information or by self-organization of interacting cells, and it can require alternative specification of cell-types from a common progenitor or sorting-out of cells that are heterogeneous already. Elucidating the mechanisms required to pattern cells in diverse tissues and organs is fundamental to understanding development and how phenotypes evolve.

The alternating dark (blue) and light (yellow) pigmented stripe pattern of adult zebrafish *Danio rerio* (**Fig. 1a)** is a useful model for dissecting patterning mechanisms^4,5,7,11,12^. Cells within the dark stripes include black pigment-containing melanophores; cells in the light stripes (known as “interstripes”) include orange-pigment containing xanthophores; and both dark stripes and light interstripes contain specialized cells called iridophores^13,14^. Iridophores are the major players for skin pattern establishment and reiteration in zebrafish. They behave as reflective cells, exhibiting angular-dependent changes in hue—iridescence—owing to membrane-bound reflecting platelets of crystalline guanine^14-16^. In the light interstripes, iridophores have a cuboidal shape and form an epithelial-like mat, presenting a “dense” morphological arrangement (**Fig**.**1b**, fluorescence panel). In the dark stripes, by contrast, iridophores are sparse in number and stellate in shape, and are sometimes referred to as having a “loose” morphology^8^ (**Fig**.**1b**, fluorescence panel). The iridophore’s importance in skin patterning has been demonstrated in experiments showing that genetically or experimentally induced deficiencies in iridophores cause pattern defects, including alterations in primary stripe positioning and boundary formation, and also lead to reductions or losses of secondary interstripes and stripes^17-21^. In addition, an evolutionary truncation in iridophore development leads to an attenuated stripe pattern in the zebrafish relative *D. nigrofasciatus*^*22*^.

**Figure 1.**
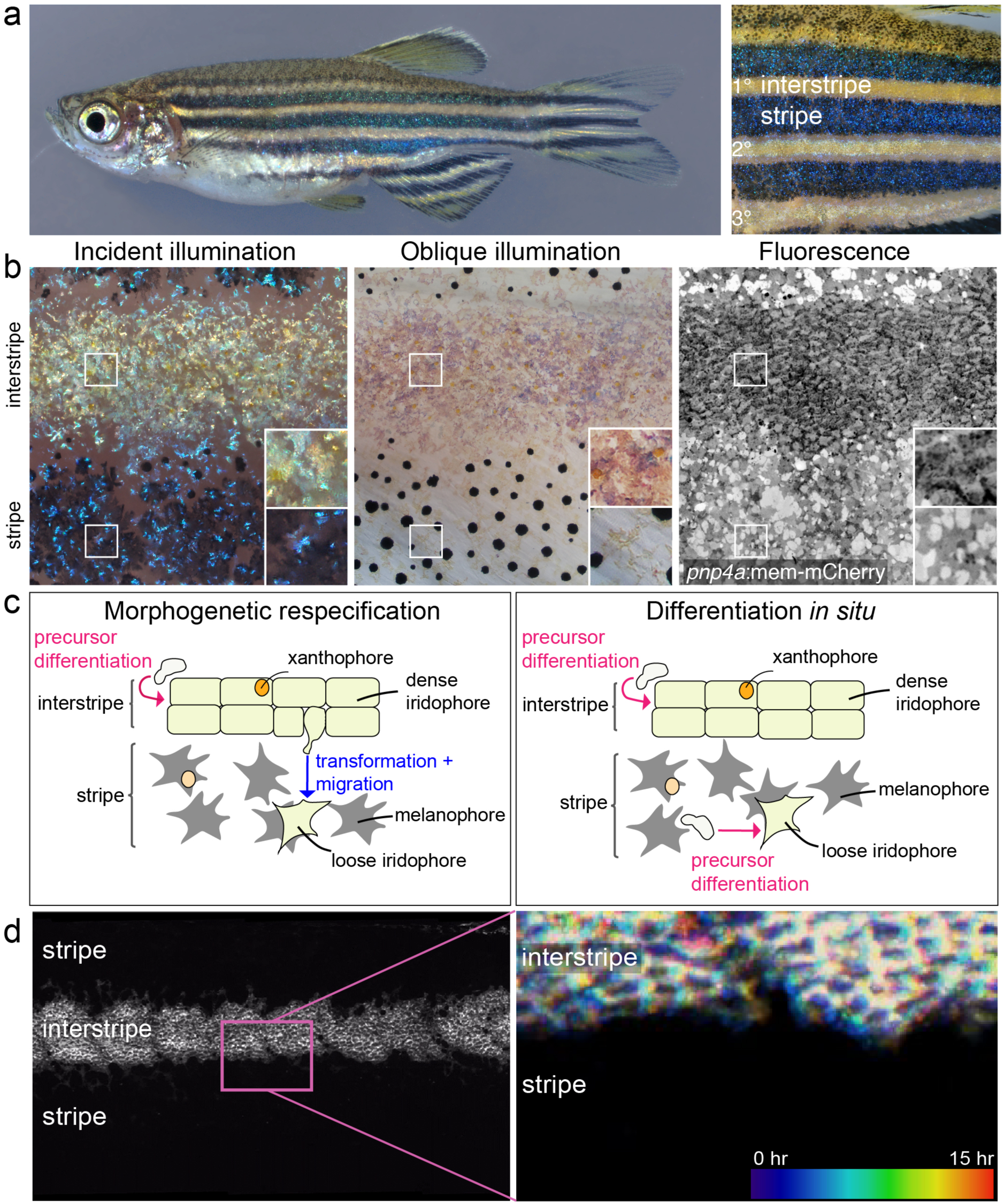
Anatomy, development and models of zebrafish adult pigment patterning. **(a)** Left panel, an adult zebrafish showing light interstripes with intervening dark stripes. Right panel, a closeup showing the primary interstripe (1°) which develops first with stripes above and below, followed by secondary interstripes ventrally (2°) and dorsally with additional stripes, and ultimately a tertiary (3°) interstripe and stripe. **(b)** Closeups of first-forming (primary) interstripe and stripes, illustrating overall pattern features as well as morphologies and arrangements of iridophores. All panels are the same location in a single animal. Left panel is incident illumination showing iridescence of iridophore-reflecting platelets with yellowish tinge in the interstripe and bluish tinge in the stripe. Center panel is oblique illumination revealing surface features and non-iridescent colors of iridophores. Right panel is membrane-targeted mCherry (mem-mCherry) driven at high levels in iridophores by regulatory elements of *purine nucleoside phosphorylase 4a* (*pnp4a*) (Eom et al., 2015^45^; Saunders et al., 2019^25^; Spiewak et al., 2018^22^) to reveal cell boundaries and arrangements. Pixel values are inverted for easier comparison to brightfield images. **(c)** Two models for iridophore patterning in interstripes and stripes. In the morphogenetic respecification model (left panel), initially densely-packed, cuboidal iridophores begin adopting a loose morphology as they and their progeny migrate out to populate the prospective stripe. In the differentiation *in situ* model (right panel), iridophores residing in interstripes and stripes are different cell types that have differentiated ‘in place’ from a precursor population. Hence, loose iridophores in stripes are not lineally related to dense iridophores in interstripes. **(d)** The flank of a 7.5 standardized standard length (SSL) *pnp4a*: mem-mCherry fish. Left panel; fluorescence image showing the arrangement of labeled cells in the dense primary interstripe. Right panel; pseudo-temporal coloring representation of a 15 hour time-lapse movie (zoomed to the region outlined in ‘d’) revealing that interstripe iridophores migrate primarily in the anteroposterior direction, with no apparent dorsoventral migration into the stripe region.

An elegant model explaining the iridophore’s role in stripe and interstripe formation links pattern establishment and reiteration to changes in iridophore morphology, proliferation and migration^7-9^ (**Fig. 1c**, left panel). Densely arranged iridophores are proposed to first proliferate to fill the primary interstripe. Some of these cells then adopt a loose shape and migrate out into the stripe zone where they continue to proliferate. Subsequently, some loose iridophores reaggregate to adopt a dense morphology and thereby initiate secondary interstripes. The iridophore shape transitions from dense-to-loose and loose-to-dense are thought to resemble epithelial-to-mesenchymal transitions (EMT) and mesenchymal-to-epithelial transitions (MET), respectively. Signals by melanophores and xanthophores are proposed to determine the specific morphologies adopted by iridophores. Consistent with this idea, quantitative models incorporating proposed dynamic morphological changes of individual iridophores are able to produce stripe patterning and robustness^23,24^.

Key predictions of the above model, hereafter referred to as “morphogenetic respecification”, are that some interstripe iridophores undergo an EMT-like transformation and migrate out from the interstripe zone, while presumably maintaining attributes (e.g., reflecting platelet composition and optical properties) other than their overall shape. In testing these predictions through a variety of approaches, we found that individual iridophores did not migrate out from the interstripe into the stripe. Instead, iridophores assumed a particular morphology at the time of their differentiation according to the presence or absence of melanophores and this morphology remained fixed thereafter. We also observed that interstripe and stripe iridophores exhibited distinct organizations of guanine-reflecting platelets (i.e., crystal types) conferring intrinsic differences in color, and that only stripe-localized iridophores could modulate reflecting-platelet spacings physiologically (blue → yellow). Furthermore, interstripe and stripe iridophores had distinct transcriptomic states. Based on these results, we propose a new model for stripe pattern formation in the adult zebrafish in which iridophore precursor cells undergo “differentiation *in situ*” into distinct iridophore types (i.e., crystallotypes) (**Fig. 1c**, right panel). This process would depend on factors in the iridophore environment that impact the specification and subcellular organization of specialized organelles within iridophore-precursors.

## Results

### Time-lapse imaging reveals iridophores do not migrate out from the interstripe

Stripe pattern establishment and reiteration in the zebrafish has been proposed to occur through morphogenetic respecification, in which iridophores differentiate to form a primary interstripe and then these cells or their progeny migrate out to contribute to stripes as well as secondary interstripes and stripes^7-9,23^ (**Fig. 1c**, left panel). Individual cells would switch morphologies as appropriate to pattern context, undergoing morphogenetic respecification via processes resembling EMT or MET.

To test this model, we examined iridophore behaviors by time-lapse imaging of membrane targeted mCherry driven by regulatory elements of *pnp4a*^22,25^. If morphogenetic respecification accounts for variation in iridophore morphology and patterning, then events resembling EMT or MET should be observable: densely packed iridophores of the completed interstripe should delaminate to populate the stripe, whereas loosely arranged iridophores of completed stripes should aggregate to initiate new interstripes. In over 300 hours of recordings, we observed no instances in which interstripe iridophores—having the dense morphology—delaminated from their neighbors and assumed the loose morphology (**Fig. 1d**; 14,475 total cells, including 1,637 cells located at interstripe edges). Likewise, interstripe iridophores that divided yielded daughter cells that remained in the interstripe (981 divisions including 160 at interstripe edges; **Fig. 1d, Extended Data Fig. 1a, Supplementary Movies S1, S2**). These observations do not support the morphogenetic respecification model.

A different way to produce cells in distinct locations having distinct morphologies would be if iridophores populate interstripes and stripes by differentiating from a progenitor not yet specified to type (**Fig. 1c**, right panel). Iridophore morphology in this model would emerge by “differentiation *in situ*” in response to context-appropriate signals, and that same morphology would be retained thereafter by the cells or their progeny. A prediction of this model for iridophore patterning is that new cells should begin to express markers of iridophore differentiation during pattern formation. Consistent with this hypothesis, we frequently observed cells acquire or increase *pnp4a*:mem-mCherry expression within developing stripes (**Extended Data Figs. 1b-e, Supplementary Movies S2–S5**).

Quantitative image analyses of proliferation and migration further supported pattern development by a mechanism of differentiation *in situ*. We found that proliferation of loose iridophores within stripes was greater than dense iridophores within interstripes^22^(**Extended data Fig. 2a**). Moreover, iridophores within interstripes tended to divide along an anterior–posterior plane, consistent with the known faster growth along this axis than dorsoventrally^26^ (**Extended Data Figs. 1b, 2b**). By contrast, planes of division by stripe iridophores were more uniformly distributed, in keeping with a rapid and relatively uniform occupancy of prospective stripe regions (**Extended data Figs. 1b-e, 2b)**. Similarly, movements of dense iridophores were negligible, whereas loose morphology iridophores could migrate up to several cell diameters and these movements tended to be biased away from the first interstripe (**Extended Data Figs. 2c, 2d, Supplementary Movie S5**).

### Fate-mapping and repeated imaging support a model of differentiation in situ

We next devised a further set of experiments to challenge the two models of iridophore patterning by following the long-term fates of iridophores marked by photoconversion (green → magenta) of nuclear-localizing *pnp4*:nucEos fluorescent protein. In this experiment, photoconverted “old” iridophores will acquire white-colored nuclei over time due to their having both newly synthesized nucEos^un^ (unconverted green) and photoconverted nucEos^conv^ (magenta) in their nuclei; by contrast, “new” iridophores formed from precursor cells will have only nucEos^un^ (green) in their nuclei. (**Fig. 2a**)^27,28^. We reasoned that if individual iridophores change their morphological states and migrate out to contribute to both interstripes and stripes, as predicted by the morphogenetic respecification model, then marking cells in one pattern element should later yield marked cells in both pattern elements. On the other hand, if individual iridophores are fixed for their morphological state and contribute only to interstripes or stripes, as predicted by the differentiation *in situ* model, marked cells should retain their morphology and be confined to their original pattern element.

**Figure 2.**
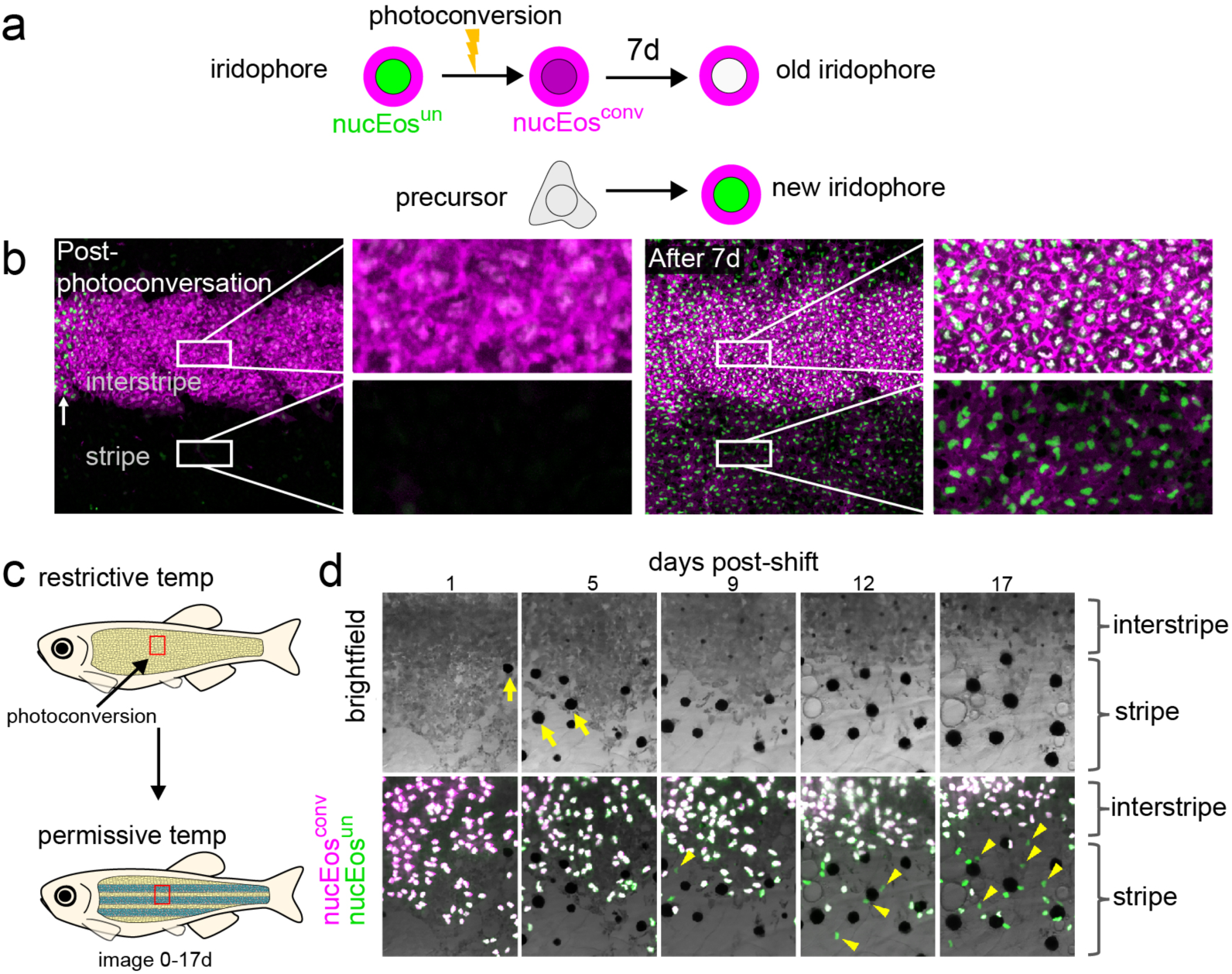
Photoconversion experiments to test models of pattern development and remodeling. **(a)** Fish were created that have iridophores expressing nuclear localizing Eos (nucEos^un^, green) and a membrane-targeted mCherry (mem-Cherry, magenta) driven by regulatory elements of *pnp4a*. Following photoconversion of an iridophore population, the converted nuclei will appear magenta (nucEos^con^). After 7 days, previously photoconverted nuclei will appear white (due to the combination of ‘new’ green proteins and ‘old’ magenta proteins), whereas nuclei of newly differentiated cells will appear green. **(b)** Tracking photoconverted iridophores in the interstripe revealed stripe iridophores do not derive from the interstripe population. Following photoconversion of a region in the primary interstripe of a fish at 7.5 SSL, all nuclei appeared magenta (Post-photoconversion), with surrounding mem-Cherry labeled plasma membrane magenta-colored as well (left panel). After seven days of additional development (8.6 SSL), at which time iridophores now populate the primary stripe, only nuclei with green signal are seen in the stripe zone, whereas interstripe nuclei are primarily white (right panel). In zoom-up images, interstripe iridophores are seen to have continued to proliferate and retained nucEos^conv^, acquiring new nucEos^un^ (making their nuclei white) (upper inset, right panel). Stripe iridophores, by contrast, lacked nucEos^conv^ and expressed only nucEos^un^ (making their nuclei green) (lower inset, right panel). **(c)** Use of a temperature-sensitive *mitfa*^*vc7*^ fish to examine the effect of conditional melanophore development on iridophore pattern remodeling. In this experimental setup, iridophores were labeled only with a nuclear-localizing Eos (nucEos^un^, green), so that following photoconversion their converted nuclei would appear magenta (nucEos^con^). **(d)** Brightfield (upper) and fluorescence superimposed on brightfield (lower) following photoconversion and shift to permissive temperature to drive onset of melanophore differentiation. Iridophores labeled by nucEos expression were photoconverted at the beginning of the experiment and followed over 17 days to distinguish newly differentiating iridophores (green nuclei) from previously differentiated iridophores (white). As melanophores differentiated (see yellow arrows in top panel), the region of dense morphology iridophores receded dorsally. This change was accompanied by differentiation of new iridophores having green nuclei (see yellow arrowheads in bottom panel) in the newly forming stripe.

Immediately after photoconverting a region in the interstripe zone, all iridophores in this region had magenta nuclei, whereas iridophores in regions not targeted for photoconversion, including a very few loose iridophores already present in the stripe zone, had only green nuclei (**Figs. 2b**, post-photoconversion). After 7 days, only iridophores in the interstripe zone had white nuclei, whereas newly-formed iridophores, having green nuclei (indicative of their acquiring *pnp4* expression), could be seen mostly in the stripe zone (**Figs. 2b**, After 7 day). The presence of white-colored nuclei in the interstripe and their absence in the stripe clearly indicates that interstripe marked cells did not migrate, favoring the model of differentiation *in situ*. Additionally, we found that the formation of secondary interstripes was characterized by the development of cells newly expressing *pnp4* within this region, suggesting differentiation with subsequent proliferation rather than active aggregation (**Extended Data Fig. 3**).

The above analyses focused on a region in the middle of the flank. Because iridophore behaviors may differ between anatomical regions, we extended our analyses by examining distributions of *pnp4*:mem-mCherry+ cells in entire, individual fish imaged daily over 33 d. These analyses also revealed extensive differentiation of iridophores without indications of EMT (**Extended Data Figs. 4** and **5a**,**b Supplementary Movie s6**). In some instances, patches of dense iridophores anteriorly appeared to split between primary interstripe and ventral secondary interstripes, perhaps owing to rapid expansion of the flank directly over the swim bladder (**Extended Data Fig. 5d**; **Supplementary Movie S7**). In a minority of larvae (∼20%), patches of 5–10 closely associated iridophores developed anteriorly, dorsal to the primary interstripe. Cells in these patches sometimes maintained their tight associations and became incorporated into the secondary dorsal interstripe. In other instances, such cells were incorporated instead into the stripe (**Extended Data Figs. 5c**,**e**). Contrary to the expectations of the morphogenetic respecification model^8,23^, the few of these cells that transitioned from a nascent dense morphology to a loose morphology occurred already within prospective stripe regions. These observations highlight subtle region-specific differences in patterning events and suggest that, had state transitions occurred in a majority of cells or over a broader anatomical area, as predicted in the morphological respecification model, they should have been observed. That they were not observed lends further support to the model of differentiation *in situ*.

### Role of melanophores in iridophore pattern remodeling

Because melanophores reside in stripe-but not interstripe-zones, we wondered whether iridophore pattern remodeling (i.e., dense-versus loose-arrangement morphology) is impacted by melanophore presence. Prior work has hinted at this possibility as mutants for melanophore-inducing trancription factor (*mitfa*), which lack melanophores, have the dense-morphology iridophores (characteristic of interstripe zones) over a broader area than in wild-type fish^18,29^. To explore this further, we used a temperature-sensitive allele, *mitfa*^*vc7*^, that allows conditional differentiation or ablation of melanophores^28,30,31^ (**Fig. 2c**), and then examined the phenotypes of iridophores in different skin areas.

Cells in a defined area were marked by nucEos photoconversion and followed over time after shifting between temperature regimes in order to assess whether iridophores in newly arising stripes, or regions newly devoid of stripes, were derived either from previously differentiated or newly differentiated cells. When fish were shifted from restrictive temperature, where they lacked melanophores, to permissive temperature, where melanophore differentiation could occur, we found that pre-existing, dense morphology iridophores receded and new iridophores differentiated into a loose arrangement in regions where melanophore differentiation had occurred (**Fig. 2d**; **Extended Data Fig. 6a**). Reciprocal temperature shifts to ablate melanophores led to a similar loss of pre-existing loose iridophores (through population turnover) and the differentiation of new dense iridophores (**Extended Data Fig. 6b**). Though we cannot exclude the possibility that some pre-existing iridophores were incorporated into remodeled pattern elements, these results suggest that the presence of melanophores has a major effect on the pattern remodeling of iridophores, specifically, in promoting a loose morphology.

### Distinct crystal morphology and ultrastructural organization but shared chemistry of loose and dense iridophores

Differences in iridophore morphologies (dense/cuboidal vs loose/stellate) and our failure to observe transitions between these two states, raised the possibility that iridophores of dense/cuboidal morphology in interstripes and loose/stellate morphology in stripes represent distinct cell subtypes, analogous to neuronal subtypes^32^. To test this possibility, we evaluated the subcellular architecture, physiology and gene expression of dense/cuboidal iridophores in stripes versus loose loose/stellate iridophores in interstripes.

Because iridophores depend for their iridescence on stacks of membrane-bound reflecting platelets consisting of crystalline guanine^5,16^, we first asked whether numbers, sizes or arrangements of these crystals differ between iridophores found in interstripe versus stripe regions. To visualize guanine crystals *in situ* required a reagent that would adhere to guanine crystals and so screened 12 cell-permeable dyes, chosen for their ability to form both hydrogen bonds and pi-stacking interactions. We found that Malachite Green efficiently bound guanine and therefore used it to examine guanine crystal organization in iridophores from interstripe versus stripe regions. Incident illumination images of the stripe zone showed blue, loosely distributed iridophores on top of black melanophores, whereas images of the interstripe zone showed dense silvery iridophores covered by yellow xanthophores (**Fig. 3a**,**b**, incident illumination). Notably, Malachite Green labeling of guanine crystals within iridophores in these two zones revealed tightly stacked arrays of crystals in loose iridophores from the stripe but markedly disordered arrays of crystals in dense iridophores from the interstripe (**Fig. 3a-b**, upper panels).

**Figure 3.**
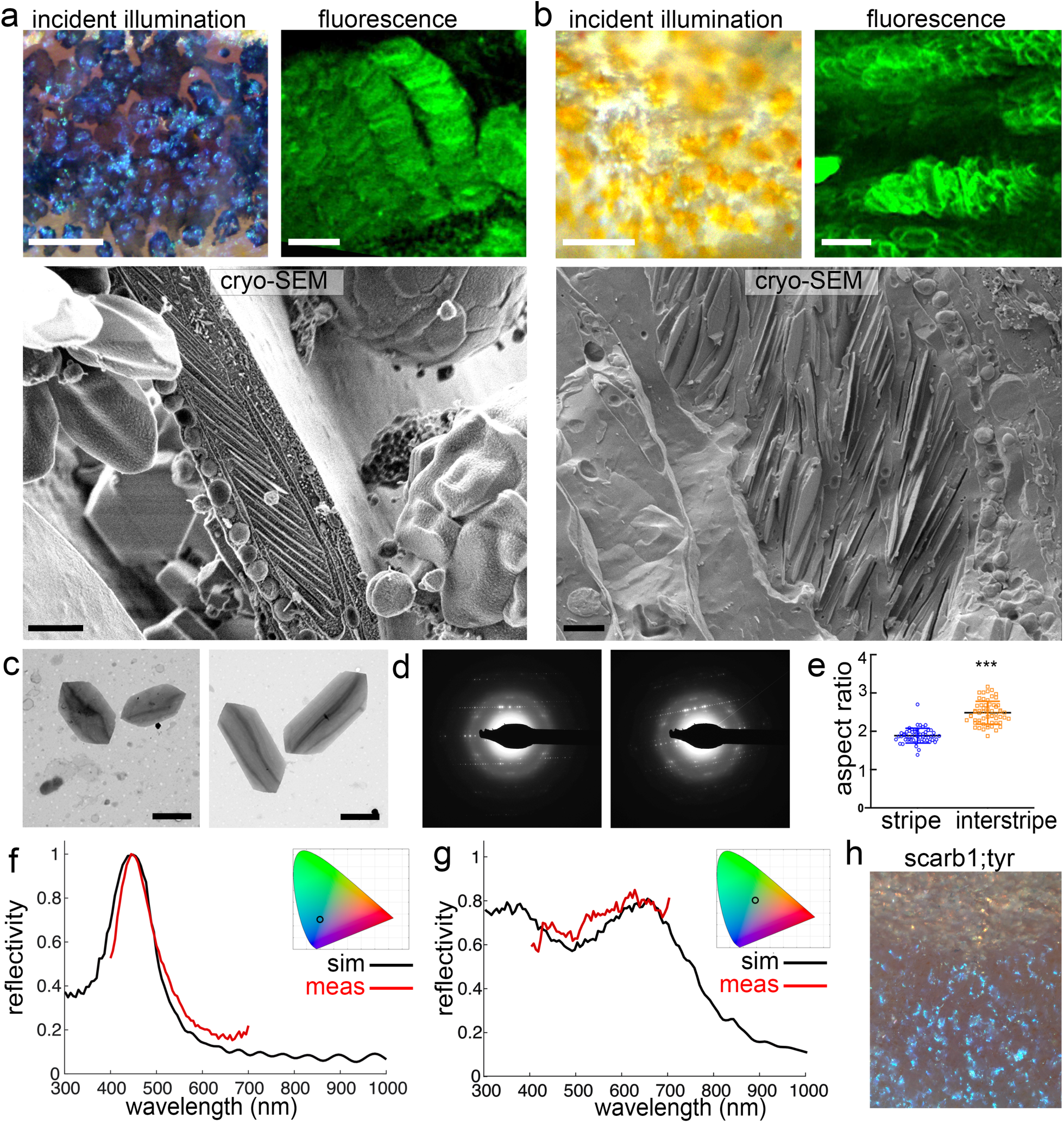
Loose-versus dense-iridophores have distinct crystal morphologies and ultrastructural organizations but shared chemistry. **(a)** Loose iridophores in stripe region viewed by incident illumination, fluorescence and high pressure–frozen, freeze-fractured cryo-SEM (Cryo-SEM). The incident illumination image shows blue iridophores on top of black melanophores; the fluorescent image reveals malachite green (MG) labeled iridophores with highly ordered arrays of guanine crystals; and the Cryo-SEM image shows iridophore cytoplasm with highly disordered arrays of crystals. **(b)** Dense iridophores in interstripe region viewed by incident illumination, fluorescence and Cryo-SEM. The incident illumination image shows silvery iridophores covered by yellow xanthophores; the fluorescent image reveals MG labeled iridophores with disordered arrays of guanine crystals; and the high pressure–frozen, freeze-fractured cryo-SEM micrograph shows iridophore cytoplasm with disordered arrangements of crystals. **(c-e)** TEM analysis of crystals isolated from iridophores from either the stripe or the interstripe regions. (c), TEM micrographs of crystals isolated from stripe (left panel) and interstripe regions (right panel); (d), TEM-based electron diffraction of crystals isolated from stripe iridophores (left panel) and interstripe iridophores (right panel); (e) graph of aspect ratio (length/width) of stripe iridophores (blue) and interstripe iridophores (orange), *p*<0.0001. **(f)** Simulated reflection (black) and measured reflection (red) from a stripe iridophore. **(g)** Simulated reflection (black) and measured reflection (red) from an interstripe iridophore. Insets in both (f) and (g) show the corresponding reflectance color on a CIE (International Commission on Illumination) chromaticity space diagram. **(h)** Incident illumination image of an adult fish lacking melanin in melanophores and carotenoids in xanthophores due to mutations in *tyrosinase* and *scarb1*, respectively. The image shows iridophore-type specific coloration is independent of melanin and carotenoids, consistent with reflectance data obtained for stripe (f) and interstipe iridophores (g).

To assess ultrastructural organization of iridophores under near-physiological conditions, in which crystal organization and cytoplasmic spacing are likely to be retained, we used cryogenic scanning electron microscopy (cryo-SEM). Crystal arrays of loose iridophores from stripes were remarkably ordered, with 20 to 30 layers of parallel crystals having an average thickness of 27±7 nm (*n*=82), neatly separated by thin layers of cytoplasm of average thickness 131±24 nm (*n*=91) (**Fig. 3a**, lower panel). By contrast, crystal arrays of dense iridophores from interstripes were disordered, varying in both orientations and spacings between crystals (**Fig. 3b**, lower panel), with 30–40 crystals per cell, and a similar average crystal thickness of 25±8 nm (*n*=130) and an average cytoplasm spacing of 186±81 (*n*=145).

Beyond differences in crystal arrangements, the shapes and sizes of crystals appeared to differ between loose iridophores in stripes and dense iridophores in interstripes. To quantify these differences, we isolated skin separately from stripes (**Fig. 3c**, left panel) and interstripes (**Fig. 3c**, right panel) and extracted crystals for transmission electron microscopy (TEM) and electron diffraction (ED) analyses. While the crystals in cells from both tissue regions comprised plates of β-guanine (**Fig. 3d**), crystals isolated from stripe iridophores were smaller (3.9±0.4 µm vs. 5.3 ±0.9 µm) and had smaller aspect ratios than crystals from interstripe iridophores (1.9±0.2 µm vs. 2.5 ±0.3 µm; n=57, 60) (**Fig. 3e**). *In situ* Raman spectroscopy of individual cells further validated that crystals in loose iridophores in stripe zones and dense iridophores in interstripe zones consist of β-guanine, and failed, within the accuracy afforded by these measurements, to reveal other components, suggesting that differences in crystal morphology are not related to their chemistry (**Extended Data Fig. 7**).

### Iridophore subtypes differ in their optical properties

Differences in colors reflected by stripe iridophores (blue) and interstripe iridophores (silvery-yellow) have sometimes been ascribed to influences of pigments contained within melanophores and xanthophores, respectively. Given the differences we observed in reflecting platelet architectures of iridophores from stripes versus interstripes, however, we reasoned that reflected spectra might be intrinsic properties of iridophore subtypes. Consistent with this hypothesis, we found close matches between spectra predicted from simulations based on a Monte Carlo transfer matrix^33^ (with morphometic data derived from cryo-SEM) and empirical reflectance spectra recorded for individual cells by hyperspectral imaging microscopy^34^ (**Fig. 3f, g**). Indeed, simulations for ordered-crystal iridophores from stripes predicted a peak in the blue region at 450 nm approaching unity reflection, whereas simulations for disordered-crystal iridophores from interstripes, predicted a broad wavelength reflection. In addition, while the reflection from the ordered-crystal iridophores was highly dependent on the angle of incident light, the reflection from disordered-crystal iridophores was not (**Extended Data Fig. 8)**.

We further found that intrinsic differences in iridophore optical properties could generate a strong contrast in color between stripes and interstripes independent of pigments in other cell types. This was manifested in fish that lack both melanin in melanophores and carotenoids in xanthophores, owing to mutations in both *tyrosinase* and *scarb1*, respectively^25^. Here, differences in color between stripes and interstripes (i.e., blue versus silvery-yellow) persisted even in the absence of other pigments (**Fig. 3h**). Together, these results demonstrate the intrinsic differences in optical properties between ordered-crystal iridophores of stripes and disordered-crystal iridophores from interstripes.

### Disordered-crystal and ordered-crystal containing iridophores remain distinct throughout development

To map the structural organization of iridophores across the entire skin pattern, we used synchrotron-based micro X-ray diffraction, which allows large areas to be scanned while still providing information on orientations and anisotropy of crystal arrays at the level of a single cell. In this system, high angular distribution diffractions having a full-ring signal are indicative of crystal orientations that vary (i.e., disordered), whereas low angular distribution diffractions having a punctuated-ring signal are indicative of crystals that are consistently oriented (i.e., ordered)^15^. Dorso-ventral line scanning across the flank of the fish demonstrated there were consistent differences in structural organization of stripe vs. interstripe regions (**Fig. 4a**). Specifically, based on their (012) and (002) diffraction planes^15,35^, crystal plates in iridophores of the stripe zone were well oriented (i.e., ordered) (**Figs. 4a**, panels 1 and 3), whereas those in the interstripe zone were non-aligned (i.e., disordered) (**Figs. 4a**, panels 2 and 4).

**Figure 4.**
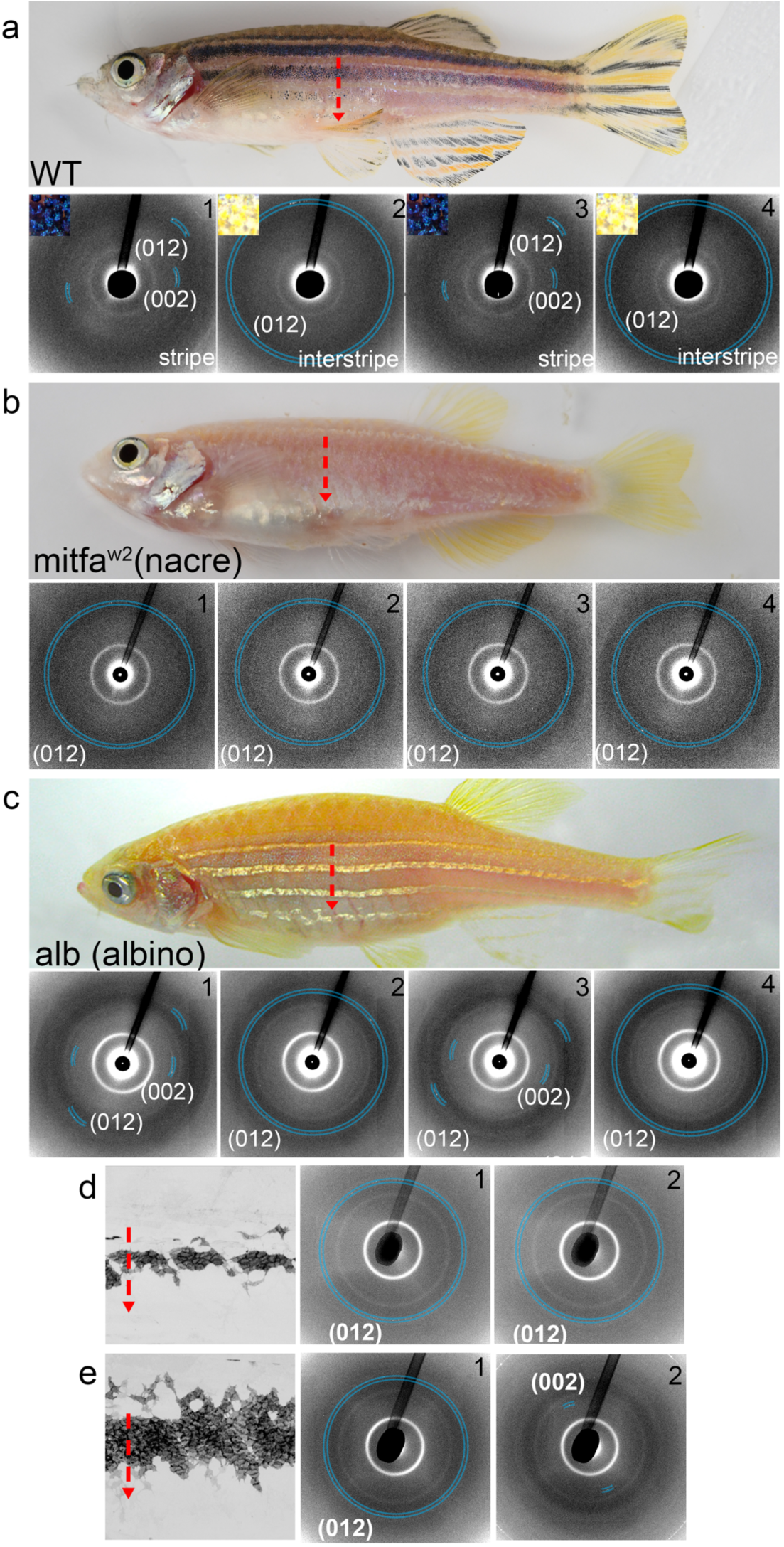
Disordered- and ordered-crystal-containing iridophores remain distinct throughout development. **(a)** Upper panel shows wildtype zebrafish with the red vertical dotted arrow showing where X-ray diffraction measurements were made. Lower panels 1 through 4 show X-ray diffraction pattern measurements in stripe and interstripe regions, with upper left insets showing the incident illumination differences in these regions. Diffraction patterns collected in the stripe regions (1 and 3) had low angular distributions with a punctuated-like signal, indicating iridophore crystals in these regions are parallel to one another. Diffraction patterns collected in the interstripe regions (2 and 4), by contrast, exhibited high angular distributions with a full-ring signal, indicating iridophore crystals in these regions are not well aligned. **(b)** X-ray diffraction measurements as in (a) made in Mitfa-w2 mutant (nacre) fish using a vertical line scan across the trunk of the fish. The typical diffraction pattern of the ordered stripe iridophore is missing in this line scan, and the observed diffractions are of high angular distribution (“full ring). **(c)** X-ray diffraction measurements as in (a) made *albino* mutant (*alb*). The overall diffraction pattern resembles that of wild type fish, with highly ordered diffraction patterns of the (002) and (012) diffraction planes throughout the stripe region (1 and 3), and high angular distribution of only the (012) diffraction plane throughout the interstripe region (2 and 4). **(d)** X-ray diffraction patterns from a vertical line measured across the trunk of a ∼6 SSL wild-type zebrafish. Panels 1 and 2 show X-ray diffraction patterns from areas in the 1° interstipe and adjacent to the 1° interstipe, respectively. Both patterns show a high angular distribution of the (012) diffraction plane. (**e)** X-ray diffraction patterns from a vertical line measured across the trunk of a ∼6.9 SSL wild-type zebrafish. Panels 1 and 2 show X-ray diffraction patterns from areas in the 1° interstipe and adjacent to the 1° interstiperespectively. A low angular distribution diffraction of the (002) plane (2) is visible just adjacent to the first interstripe region (1).

Our *mitfa*^*vc7*^ photoconversion results (see **Fig. 2c**) raised the possibility that melanophores promote the differentiation of progenitors into iridophores with ordered crystal arrays. We tested this idea using micro X-ray diffraction to evaluate the crystals architecture in iridophores of two zebrafish mutants: null-allele *mitfa*^*w2*^ and *albino*. In *mitfa*^*w2*^ mutants, melanophores are missing owing to a defect in their specification; in *albino* mutants, melanophores are present but lack melanin^29,36^. We reasoned that if melanophores drive iridophore differentiation towards the ordered crystallotype, then *mitfa*^*w2*^ mutants should be deficient in iridophores having ordered crystals, whereas *albino* mutants should retain ordered iridophores, similar to the wild type. Line scans across the flanks of *mitfa*^*w2*^ fish revealed mostly high angular distribution (012) diffractions, typical of the disordered crystallotype (**Fig. 4b**). Scanning the entire fish, showed some diffraction patterns corresponding to ordered iridophores, but these were located towards the posterior and were a minor component of the diffractions (**Extended Data Fig. 9**). The same analysis on *albino* fish revealed alternating diffraction patterns similar to that seen in wild-type (**Fig. 4c**). These results suggest that melanophores enhance the differentiation of ordered crystallotype iridophores.

We next examined the relative developmental timing of precursor differentiation into disordered and ordered crystallotypes by assessing micro X-ray diffraction patterns over ontogeny. In fish of 6.0 mm standardized standard length (SSL) and 6.5 SSL, which have only a single interstripe and very few adult melanophores^26^, we observed only disordered crystallotype iridophores (having high angular distribution diffraction patterns of the (012) plane) (**Fig. 4d**; **Extended Data Fig. 10**). When fish were ∼6.9 SSL, with a substantial complement of melanophores and loose iridophores, low angular distribution diffraction patterns of the (002) plane, typical of ordered crystallotype iridophores, became visible (**Fig. 4e**). These results supported the idea that precursor cells differentiate into ordered crystallotype iridophores only after the differentiation of disordered-crystallotype iridophores and in the presence of melanophores.

### Ordered and disordered crystallotypes exhibit distinct transcriptomic signatures

We next compared the transcriptomic signatures of ordered and disordered crystallotype iridophores (i.e., from stripes and interstripes) by single cell RNA-sequencing (scRNA-seq) (**Fig. 5a**). Dimensionality reduction followed by unsupervised clustering revealed six clusters, three of which (clusters 4–6) contained cells expressing known markers of iridophores^37^ (i.e., *tfec, gpnmb*) (**Fig. 5b**). Expression of purine synthesis pathway genes, markedly upregulated^25^ in iridophores, further validated this initial assignment (**Extended Data Fig. 11**). Next, we tested whether iridophore clusters recovered by scRNA-seq corresponded to their anatomical sites of origin. Supporting this possibility, 98% of cells in cluster 5 originated from interstripes, and 85% of cells in cluster 4 originated from stripes (**Fig. 5c**). Cells of cluster 6 were split between interstripe (63%) and stripe (37%) (**Extended Data Fig. 12**). Pseudotemporal ordering showed that ordered-crystal iridophores and disordered-crystal iridophores were associated with different branches in trajectories of inferred differentiation, whereas cluster 6 iridophores spanned across branches (**Fig. 5d**). Several hundred loci were differentially expressed between cells of clusters 4 and 5 (**Fig. 5e** and **Supporting information**), suggesting candidate genes that may contribute to structural or other differences between stripe and interstripe iridophores. We concluded from these analyses that iridophores of interstripe and stripe zones are transcriptionally distinct.

**Figure 5.**
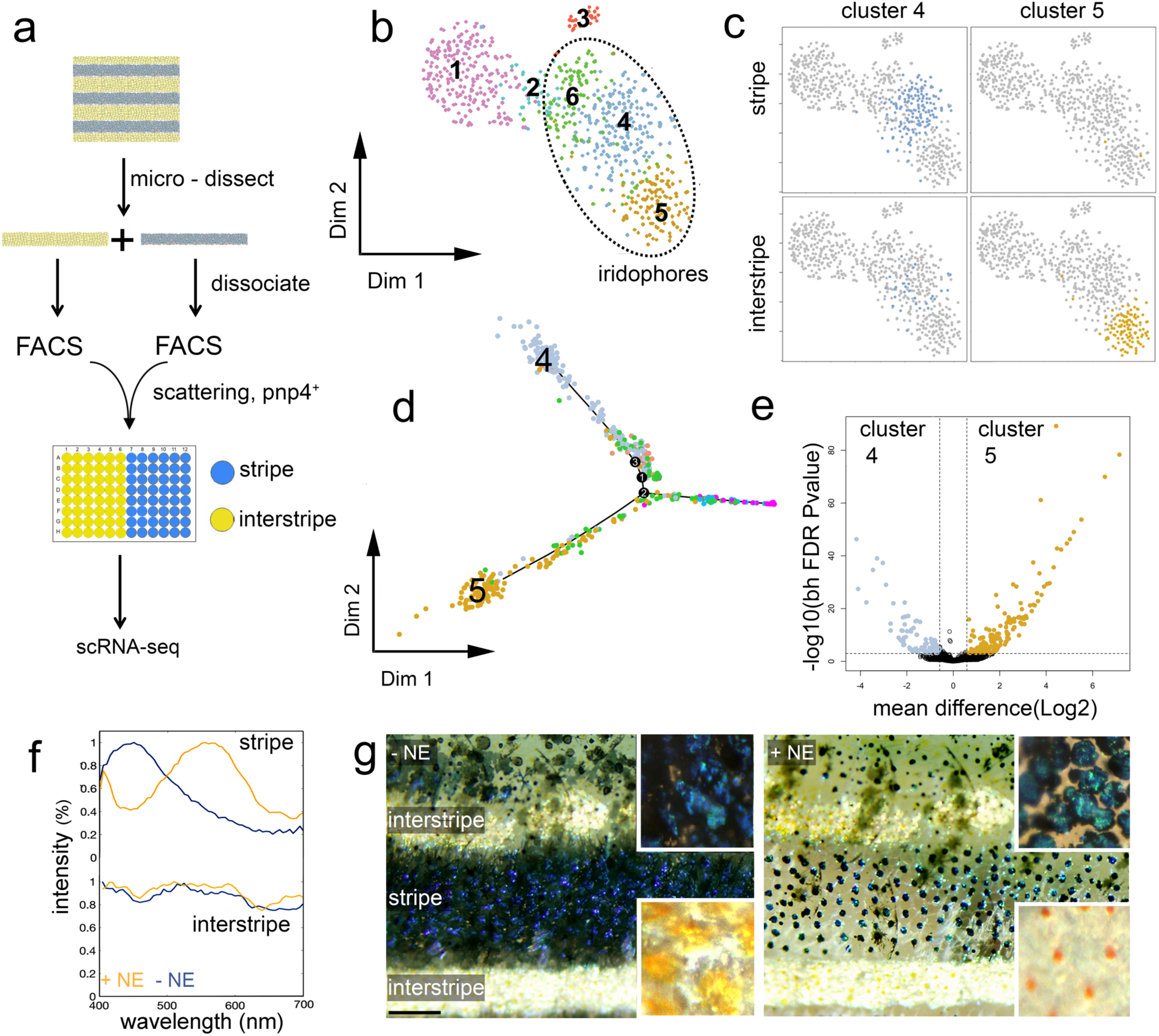
Disordered- and ordered-crystal-containing iridophores exhibit distinct transcriptomic signatures and response to stimuli. **(a)** Experimental design of single cell RNA sequencing experiment. **(b)** Two-dimensional tSNE representation of the collected skin cell clusters (dashed ellipse marks iridophores). **(c)** Anatomical origin of iridophores clusters. **(d)** Pseudo-time trajectory of the collected skin cells. **(e)** A volcano representation of differentially expressed genes between clusters 4 (light blue) and 5 (orange). **(f-g)** The response of an adult zebrafish skin pattern to norepinephrine (NE) stimuli. (f) is the optical response of individual iridophores from the stripe (upper panel) and interstripe (lower panel). In the stripe, the reflection peak of an ordered iridophore shifts from ∼450 nm to ∼570 nm upon NE treatment. In the interstripe, only minor changes in the reflection spectra occur in response to NE. (g) is the optical response of the relaxed, untreated fish (-NE) compared to the treated fish (+NE). Note the contrast between the blue stripe and the two-flanking yellow interstripes and how this changed with NE treatment. Before treatment, a deep blue color for the stripe region and a golden-yellow color for the interstripe region is observed. After NE treatment, the contrast between the stripe and the interstripe is drastically reduced. This color change arises because NE causes melanophores in the stripe to aggregate and blue iridophore reflectance to shift from a dark-blue to green-yellow hue (see upper insets), while in the interstripe, NE causes xanthophores to aggregate and silvery iridophores to have only a minor color change (lower insets).

### Physiological color change differs between iridophore subtypes

Color pattern can be influenced physiologically^38,39^, as some types of pigment cells disperse or contract pigment granules in response to endocrine and neuroendocrine factors, including norephinephrine (NE)^40-42^. We wondered whether differential responses from the iridophore subtypes might contribute to this response. To test this, we bathed isolated fish skin in 10 µM NE solution. Upon NE treatment, interstripe-derived iridophores exhibited a ∼120 nm shift in peak reflection, whereas stripe-derived iridophores exhibited only a minor change in their overall organization of crystals and spectrum of reflected light measured by hyperspectral imaging (**Fig. 5f**). Viewing the response to NE in the context of the whole tissue also revealed a preferential color change in striped-versus interstripe-zones (**Fig. 5g**). These results support prior work showing that iridophores of different shapes respond differently to NE^40^, and further demonstrate that only stripe iridophores change their color upon NE treatment. This differential response, in conjunction with the known aggregation of granules within melanophores and xanthophores^41,42^, could contribute to the dramatic reduction in contrast between stripe and interstripe zones under NE treatment, which causes the prominent zebrafish stripe pattern to diminish (**Supplementary movie 8**).

## Discussion

Pigmentation of teleost fish has become a valuable system for understanding pattern formation in animals, including how changes in pattern-forming mechanisms lead to phenotypic variation within and between species^4,7,11^. In zebrafish, a widely accepted model links dynamic changes in iridophore shape—between a dense morphology in interstripes and a loose morphology in stripes—to establishment and reiteration of pattern^8,9,23^. In this model, iridophores of interstripes and stripes are similar cells that have merely adopted different morphologies as they migrate into different regions. Our findings of substantial differences in ultrastructure, physiology and transcriptomic state of iridophore subtypes—and the absence of morphological transitions predicted for individual iridophores—support an alternative model of differentiation *in situ* for how the reiterated stripe pattern of zebrafish develops. In this model, iridophore precursors in developing stripes and interstripes differentiate *in situ* into distinct iridophore crystallotypes with different subcellular organization and physiological responsiveness based on their micro-environment.

Strongly supporting a model of differentiation *in situ* was our finding of substantial physiological disparities between iridophores of stripes and interstripes. In particular, iridophores of interstripes had larger reflecting crystal platelets that were disordered, whereas those of stripes had smaller crystal platelets that were uniformly stacked and oriented. The colors of these cells differed as well: iridophores in interstripes were silvery-yellowish, and iridophores in stripes were blue, and these differences were autonomous properties of the cells, not a consequence of pigments contained in other pigment cells with which iridophores associate. Physiological responses also differed: disordered crystal platelets of interstripe iridophores were refractory to NE, whereas, ordered crystal platelets of stripe iridophores changed their cellular organization upon NE treatment. Finally, iridophores from interstripes and stripes had distinct gene expression profiles.

We found no evidence for another key prediction of the prior model, namely that individual iridophores should undergo state transitions as cells originating in one pattern element disperse to populate another. Photo-labeled iridophores observed over short or long periods failed to migrate between interstripe and stripe zones, even when they were challenged to undergo transitions in the context of pattern remodeling (stimulated by changes in melanophore abundance). Instead, development and remodeling of interstripes and stripes involved the *in situ* differentiation, subsequent proliferation, and in some cases migration of iridophores with morphologies appropriate to their location within the pattern. Only in a minority of fish, and in a small anatomical region, did we observe patches of initially dense iridophores assume a loose arrangement. Such behaviors occurred within prospective stripes, rather than at boundaries between interstripes and stripes, as previously postulated, and involved cells that had not yet fully differentiated.

These data all point to a model of stripe patterning that depends on differentiation *in situ*. In this model, latent progenitors associated with the peripheral nervous system that have transited to the skin during the larva-to-adult transformation expand clonally as iridoblasts—not specified to subtype—and subsequently differentiate according to cues in the microenvironment they encounter. Our data, together with those of others, suggest that some of these signals depend on melanophores^18,19,27,43^, promoting differentiation of iridoblasts towards a state having ordered reflecting crystal platelets that are physiologically responsive (blue ⟷ yellow) in stripes, and away from an alternative state of disordered crystal platelets lacking physiological responsiveness (silvery-yellow) in interstripes. Additional signals from xanthophores, iridophores, and other cell types likely contribute as well. Whether these events of specification unfold as iridoblasts expand their territory within the plane of the skin hypodermis, or as they arrive at the hypodermis after migrating from progenitor niches within the peripheral nervous system, remains to be determined. Whichever mode of iridoblast morphogenesis holds true, our findings highlight the importance of extrinsic factors that specify and promote the *in situ* differentiation of iridophore subtypes during pattern establishment and reiteration. The resulting iridophore subtypes likely allow the zebrafish to alter its skin patterning to make it more or less distinctive, a trait crucial for the fish to be able to join shoals or obscure itself^1,44^.

## Acknowledgements

We thank Gil Levkowitz and Dan Oron, for their help and advice, Ivo Zizak for his help with the diffraction measurements at the μ-Spot beamline in BESSY II. We thank Kathy Schaefer, Lihua Wang and Andrew Lemire, for their help with FACS, and scRNA-seq. We thank Rick Huang and Zhiheng Yu, for their help with ED. We thank HZB for the allocation of synchrotron radiation beamtime. We thank members of the Lippincott-Schwartz lab for their support. This work was supported by Howard Hughes Medical Institute to JLS and NIH R35 GM122471 to DMP. EJB was additionally supported by NIH T32 HD007183, DG was additionally supported by the Human Frontiers Cross-Disciplinary Postdoctoral Fellowship LT000767/2018, the European Molecular Biology Organization Long Term Fellowship and the Rothchild Fellowship. Hyperspectral analyses, MCA and DDD were supported by a Multidisciplinary University Research Initiative (MURI) - Melanin Grant, AFOSR FA9550-18-1-0142 (to DDD).

## Author contributions

Author contributions: D.G, E.J.B, D.D.D, J.C.L D.M.P., and J.L.-S. designed research; D.G, E.J.B, A.J.A, A.P, J.D.F, M.C.A, performed research; D.G, E.J.B, K.J, J.D.F, D.D.D, J.C.L D.M.P., and J.L.-S. analyzed data; and D.G, E.J.B, D.M.P., and J.L.-S. wrote the paper.

