## Supporting information for "*In situ* differentiation of iridophore crystallotypes underlies zebrafish stripe patterning"

**Supporting Information contains:**

- **Materials and Methods**
- **Extended Data Figures**
- **Supplementary Movies Legends**

**Materials and Methods –**

**Fish stocks, rearing conditions, transgenic line production, and CRISPR/Cas9 mutagenesis**

We reared fish under standard conditions (14L:10D at ~28˚C) and staging followed (Parichy, et al., 2009). Stocks were wild-type WT(AB)b, a derivative of AB*, Tg(*pnp4a*-memCherry) (Spiewak et al., Saunders et al.), temperature-sensitive *mitfa*$^{vc7}$ (Johnson et al., 2011) raised at permissive temperatures (24 °C) or restrictive temperature (33 °C), *mitfa*$^{vc7}$Tg(*pnp4a*-memCherry), *scarb1* (Saunders, et al., 2019). To generate Tg(*pnp4a*-nucEos), we used restriction enzyme cloning to replace the fluorophore in the *pnp4a*-memCherry construct with nuclear localized Eos. We injected this construct into single-cell embryos along with *Tol2* transposase mRNA using standard methods (Suster et al.) and screened for germline incorporation. CRISPR/Cas9 mutants *scarb1*;F0(*tyrp1b*-CRISPR) were generated as described in (Saunders, et al., 2019).

**Imaging**

We acquired images on Zeiss AxioObserver inverted microscopes equipped with Axiocam HR or Axiocam 506 color cameras, a Yokogawa laser spinning disk with an Evolve or Hamamatsu ORCA-Flash4.0 camera, an AxioZoom v16 stereomicroscope with Axiocam 506 color camera, or LSM800 scanning laser confocal with Airyscan detection all running ZEN blue software. For repeated daily imaging, we raised individuals in separate beakers. They were anesthetized briefly, imaged, and allowed to recover.

**Photoconversion**

We photoconverted fish expressing *pnp4a*:nucEos as a stable transgenic line (Fig 3A, S2) or mosaic (Fig 3C, S4E, S5) using the Zeiss LSM 800 scanning laser confocal with a 405nm laser and ZEN blue software. When we photoconverted the whole flank (Fig S2), we placed fish in a box lined with aluminum foil and exposed to the full intensity output from HXP 120V compact light source.

**Time lapse imaging and analyses**

Ex vivo pigment cells were imaged in their native environment on a Zeiss AxioObserver inverted microscope with a Yokogawa laser spinning disk and an Evolve or Hamamatsu ORCA-Flash4.0 camera (Eom et al., 2015). We imaged the entire flank region between the gut and caudal fin peduncle of Tg(*pnp4a*-memCherry) fish every 5 minutes for 15 hours, unless otherwise noted. Analyses focused on the area including the primary interstripe, ventral primary stripe, and ventral secondary interstripe (where applicable). All iridophores in these regions were included in proliferation and migration analyses. Proliferating iridophores were evident as single cells that rounded-up and then divided to generate adjacent daughter cells. We measured the plane of proliferation using the angle tool in ImageJ from the center of each daughter cell in the frame immediately following cytokinesis. To analyze migration, we used ImageJ to measure the start point and end point of all stripe-associated iridophores that moved ≥ one cell diameter over the course of each time-lapse movie.  Three trunks each from fish at stages 7.5SL, 10SL, and 12SL were included in behavioral analyses. Iridophores were classified as either dense (surrounded by other iridophores, bright memCherry expression on all sides), loose (stellate, dim mem-Cherry expression), or edge (~3/4 sides touching other dense iridophores, remaining side open to stripe region). Total cells analyzed: 7.5SL; D: 1656, L: 331, E: 690. 10SL; D: 4466, L: 2235, E: 523. 12SL; D: 1657, L: 1508, E: 186.

**Temperature shift iridophore challenge**

We injected *mitfa*$^{vc7}$; Tg(*pnp4a*-memCherry) with *pnp4a*:nucEos (the same used for the stable transgenic line) and Tol2 mRNA using standard methods (Suster et al.) so that *pnp4a*:nucEos expression was mosaic. We then raised the fish at 33C (restrictive temperature) or 24C (permissive temperature) until the primary interstripe, primary stripes, and ventral secondary interstripe developed at permissive temperatures. We then photoconverted the entire *pnp4a*:nucEos+ patch as described above and shifted the fish to the opposite temperature.

**Synchrotron based micro wide X-ray diffraction.** Fish larvae were euthanized in MS222 and were immersed in a physiological buffer (PBS, 136 mOsm/kg). Fish samples or entire larvae were mounted on a lead tape between two kapton windows. In situ wide X-ray diffraction (WAXD) was obtained at the µ-Spot beamline, in the synchrotron radiation facility BESSY II, Helmholtz-Zentrum Berlin für Materialien und Energie, Berlin, Germany. Samples were mounted on a y–z scanning table and scans of the sample in areas of interest were performed. The microbeam was defined by a toroidal mirror and a pinhole of either 10 μm or 30 μm diameter close to the sample, providing a beam size of approximately $10\times10\mu m^2$ or $30\times30\mu m^2$ at the sample position. An energy of 15keV ($\lambda$ = 0.82656Å) was selected by a Mo/BC multilayer monochromator. The 2D SAXS/WAXD patterns were measured by using a MarMosaic 225 CCD-based area detector (Rayonix) placed at a sample-detector distance of 286 mm. The beam center in the detector and the sample-detector distance were calibrated using a powder X-ray diffraction pattern of synthetic guanine powder standard (Sigma Aldrich). Radial integration of the 2D scattering patterns was performed by using Fit2D. The data were normalized with respect to the primary beam monitor (ionization chamber) and corrected for background caused by pinhole and air scattering.

**Hyperspectral Imaging.** Fish were euthanized in MS222, then relevant skin portions were dissected and placed in PBS solution at room temperature. Spectral measurements were performed using a PARISS® hyperspectral imaging system (Lightform Inc.), which acquires instantaneously 380-980nm spectra from each pixel of a line of pixels when pixel size was $1.25\times1.25\ \mu m^2$. The hyperspectral imager was mounted on a Nikon Eclipse 80i microscope. Spectral calibration was performed using a MIDL® $Hg^+/Ar^+$ wavelength calibration lamp (Lightform Inc.) with accuracy better than 2nm. The cells spectra were acquired under both specular reflectance and transmittance, using a tungsten halogen light source with a Nikon NCB11 filter, and a 20x 0.50NA objective. Reflectance spectra were normalized using a standard silver mirror (Thorlabs Inc.). All spectra were smoothed with a moving average of 3 and plotted using Matlab.

**TEM.** Fish were euthanized in MS222 then relevant skin portions were dissected and placed in PBS solution at room temperature. Tissue was mechanically dissected using a scalpel and then sonicated for 10 min. The obtained suspension was centrifuged for 5 min at 4000RPM, after which the pellet was discarded, and the supernatant was kept and vortexed for 1 min. this process was repeated twice. 10 µl drop of obtained suspension was placed onto a copper slot TEM grid coated with Formvar/carbon and was allowed to settle for 5 min. After which the grid was carefully blotted dry with help of a piece of filter paper. The TEM grids were imaged in a Tecnai Spirit electron microscope (FEI, Hillsboro, OR) operating at 80kV equipped with an Ultrascan 4000 digital camera (Gatan Inc, CA).

**Raman Microspectroscopy.** Samples were prepared by sandwiching prepared tissues or crystals between quartz coverslips (Electron Microscopy Sciences, Cat. No. 72255-02) and glass microscope slides (VWR, Cat. No. 16004-422). All data were collected at RT using a home-built Raman microscope, as previously described in reference[1] Briefly, the 514-nm line of an argon-ion laser (CVI Melles Griot, 35-MAP-431-200) was passed through a clean-up filter (Semrock, LL01-514-25) and then directed into a modified inverted microscope (Olympus IX71). Excitation light (~30 mW at the sample) was directed to the sample using a dichroic mirror (Semrock, LPD01-514RU-25x36-1.1) and a 60X water-immersion objective (Olympus, UPLSAPO60XW). Spontaneous Raman Stokes scattering was collected through the same objective, filtered (Semrock, LP02-514RE-25) to remove any residual excitation light or Rayleigh scattering, and then directed into a 320-mm focal length (f/4.1 aperture) imaging spectrometer (Horiba Scientific, iHR 320) through a 400 µm pinhole (Thorlabs) and a 50-µm slit, and dispersed using a 1200 g/mm grating. Individual spectra were collected for 10–15 s acquisitions (x10–15) from 500–3700 $cm^{-1}$ with high gain enabled on a liquid nitrogen cooled, back illuminated deep-depletion CCD array (Horiba Scientific, Symphony II, 1024×256 px, 26.6 mm × 6.6 mm, 1 MHz repetition rate). Bright field images were collected using a USB 2.0 camera (iDS, UI-1220-C). Daily calibration of imaging spectrometer was done using neat cyclohexane (20 µL in a sealed capillary tube). Bandpass and accuracy were found to be <12 $cm^{-1}$ and ± 1 $cm^{-1}$, respectively. All Raman spectra were corrected by applying a baseline polynomial fit (Lab Spec 6 software).

**Isolation of Cells for Single Cell RNA-Sequencing.** Fish were euthanized in MS222 and the stripe and interstiped regions were micro-dissected from fish expressing both pnp4a:palmmCherry and pnp4a:nlsEos. Stripe and interstipe regions were enzymatically dissociated separately with Liberase (0.25 mg/ml in dPBS) at 25°C for 15 min followed by manual trituration with increasingly narrower flame polished glass pipette for 3 min at a time for three times. Cells suspensions were then filtered through a 70 µm Nylon cell strainer to obtain a single cell suspension. Liberated cells were re-suspended in 1% BSA / 5% FBS in dPBS before FACS purification. The flow cytometry experiments were performed on a BD FACSAria II SORP sorter (BD Biosciences, San Jose, CA, USA) coupled with BD FACSDiva software (BD Biosciences) for instrument operation, data acquisition, and analysis. A 637 nm and a 488 nm laser were utilized for fluorophore-excitation and a 100-µm nozzle was used to generate single droplets under 20PSI sheath pressure. The applied settings were as follows: forward light scatter (FSC) detector photomultiplier tube (PMT) gain setting = 80 V with a 1.5 neutral density filter; side light scatter (SSC) detector PMT gain setting = 90 V; FSC threshold = 10,000; PE-Texas Red (PE TX Red) channel PMT gain setting = 280 V; fluorescein isothiocyanate (FITC) channel PMT gain setting = 280 V. Sample dilution and flow rate were adjusted to optimal event recordings for 96 well plate single cell sorts (below 500 processed events per sec). The population of zebrafish skin iridophores was designated based on their FSC and SSC characteristics and back-gating on fluorescence. Control zebrafish skin samples were used to gate out the highly auto fluorescent cells among the members of this population. Fluorescently labeled from either the stripe or interstripe skin samples were sorted separately. Single cells with high mCherry and Eos expression were collected into 3uL of smart-scrb lysis buffer (0.2% Triton X-100 [Sigma], 0.1 U/ul RNAse inhibitor [NEB]) using „single cell" indexed sorting mode. Five indexed 96 well plates were collected for each skin band. After sorting, the samples were spun down at 3166 rcf at 4°C for 2 minutes in an Eppendorf tabletop 5810R centrifuge (Eppendorf AG, Hamburg, Germany) and stored at -80°C until further processing.

**scRNA-Seq Library Preparation.** Four plates of each skin band were selected for sequencing. cDNA was prepared from sorted cells as described [2], with minor modifications. Reverse transcription, PCR, purification, tagmentation, and library quantification were performed as described. Libraries from eight plates were pooled equimolar and were sequenced on a NextSeq 550 flowcell with 25 bases in read 1, 8 bases in the i7 index read, and 50 bases in read 2. phiX control library (Illumina) was spiked in at a final concentration of 15% to improve color balance in read 1.

**scRNA-Seq Analysis.** Raw counts representing the enumerated expression for 32,618 features for 784 cells was imported into R (https://www.r-project.org/) in matrix form with features in rows and cells in columns. The "SingleCellExperiment" function (https://www.bioconductor.org/packages/release/bioc/html/SingleCellExperiment.html) was then used to convert this matrix into a single cell experiment object. Features not having a raw count greater than zero for all cells were removed from this object. Where after, the "calculateQCMetrics" function supported in the in the "scater" package (https://bioconductor.org/packages/release/bioc/html/scater.html) was used to enumerate the percent total counts across all MT genes per cell. These percentages were then visually compared across all cells using the "hist" function. Based on the resulting distribution, cells were then filter removed if they had a MT percent total count greater than two percent. For cells not discarded, counts for MT genes were removed and the non-MT features filtered to keep only those having a count greater than zero for at least one cell. The total number of features and the total counts per cell were then enumerated and visually compared using the "plotColData" function. Filtering decisions were then made based on this visual and cells discarded if they did not satisfy the following criteria: Log10 total counts per cell > 4.5, Log10 total counts per cell < 5.75, Log10 total features per cell > 2.15, Log10 total features per cell < 3.25. Features for surviving cells were again filtered to keep only those having a count greater than zero for at least one cell. While, counts for surviving features were next pedestalled by 1, normalized using the "calculateCPM" function, then ultimately Log2 transformed. The "sc3_estimate_k" function supported in the "SC3" package (http://www.bioconductor.org/packages/release/bioc/html/SC3.html) was then used to estimate the number of clusters of cells present (k=6) and the "sc3" function called to generate those clusters. Cell-to-cluster membership results were then exported from R via the "sc3_export_results_xls" function, inspected as a heatmap using the "sc3_plot_consensus" function, and overlaid onto a tSNE representation using the "Rtsne" function. Subsequent analyses performed based on these clusters included the pairwise comparison of means per feature using the Welch-modified t-test and the representation of those results by volcano plot. Pseudotime analysis was also performed using the "monocle" package (https://bioconductor.org/packages/release/bioc/html/monocle.html) in conjunction with "M3Drop" package for differential feature selection (http://www.bioconductor.org/packages/release/bioc/html/M3Drop.html) and the “plot_cell_trajectory” function for visualization of results. The data discussed in this publication have been deposited in NCBI's Gene Expression Omnibus (Edgar et al., 2002) and are accessible through GEO Series accession number GSE144734 (https://www.ncbi.nlm.nih.gov/geo/query/acc.cgi?acc=GSE144734

**Cryo-scanning electron microscopy (Cryo-SEM).** Fish were euthanized in MS222 then relevant skin portions were sectioned and placed in PBS solution at room temperature. The sections were then sandwiched between two metal discs (3 mm diameter, 0.1 mm cavities) and cryo-immobilized in a high-pressure freezing device (HPM10; Bal-Tec). The frozen samples were mounted on a holder under liquid nitrogen and transferred to a freeze-fracture device (BAF60; Bal-Tec) using a vacuum cryo-transfer device (VCT 100; Bal-Tec), where they were coated with a 3-nm-thick layer of Pt/C. Samples were then observed by high-resolution SEM (Ultra 55, Zeiss) using secondary electron/backscattered electron and an in-lens detector, maintaining the frozenhydrated state by using a cryo-stage operating at a working temperature of −120°C. Measurements of crystal thickness and cytoplasm spacing were taken from the cryo-SEM micrographs.

**Malachite green staining.** Fish were euthanized in MS222 then relevant skin portions were sectioned and placed in a PBS solution containing 25-50 μM malachite green, at room temperature for 30 min before imaging.

**Reflectivity simulations.** The reflectivity spectrum was simulated based on crystal thicknesses and spacing obtained from cryo-SEM images using a Monte Carlo transfer matrix calculation, as described in detail in the supporting information of [3]. In brief, the percentage of reflectivity was calculated by averaging 500 runs, assuming normal incident light, or in the angle depends studies between 0°-70°. Each layer was characterized by two variables: $n_j$, a refractive index, and $d_j$, which is the layer thickness randomly picked from the experimental distribution. Thus, for each layer we defined the following 2 × 2 matrix:

$$m_j = \begin{pmatrix} \cos\beta_j & -\frac{i}{n_j}\sin\beta_j \\ -in_j\sin\beta_j & \cos\beta_j \end{pmatrix} \quad \text{where} \quad \beta_j = \frac{2\pi}{\lambda} n_j d_j$$

The set of k double layers was characterized by an overall reflectivity 2 × 2 matrix:

$$M_j = \prod_{j=1}^{j=2k} m_j$$

The reflectivity was extracted from the following equation:

$$R = \left| \frac{(m_{11} + m_{12}) - (m_{21} + m_{22})}{(m_{11} + m_{12}) + (m_{21} + m_{22})} \right|^2$$

The refractive index of the guanine crystal plates was taken as 1.83, which was the refractive index in the direction of the impinging light. The weak dependence of the refractive index on wavelength was neglected, assuming that all the interfaces, i.e., inside a crystal stack and between stacks, were parallel. We also assumed no correlation between the crystal spacings within a single crystal stack.

**References-**


1 Flynn, J. D. & Lee, J. C. Raman fingerprints of amyloid structures. *Chemical communications* **54**, 6983-6986 (2018).

2 Cembrowski, M. S. *et al.* Dissociable structural and functional hippocampal outputs via distinct subiculum cell classes. *Cell* **173**, 1280-1292. e1218 (2018).

3 Gur, D. *et al.* Structural basis for the brilliant colors of the Sapphirinid copepods. *J. Amer. Chem. Soc.* **137**, 8408-8411, doi:10.1021/jacs.5b05289 (2015).

**Extended Data Figures-**

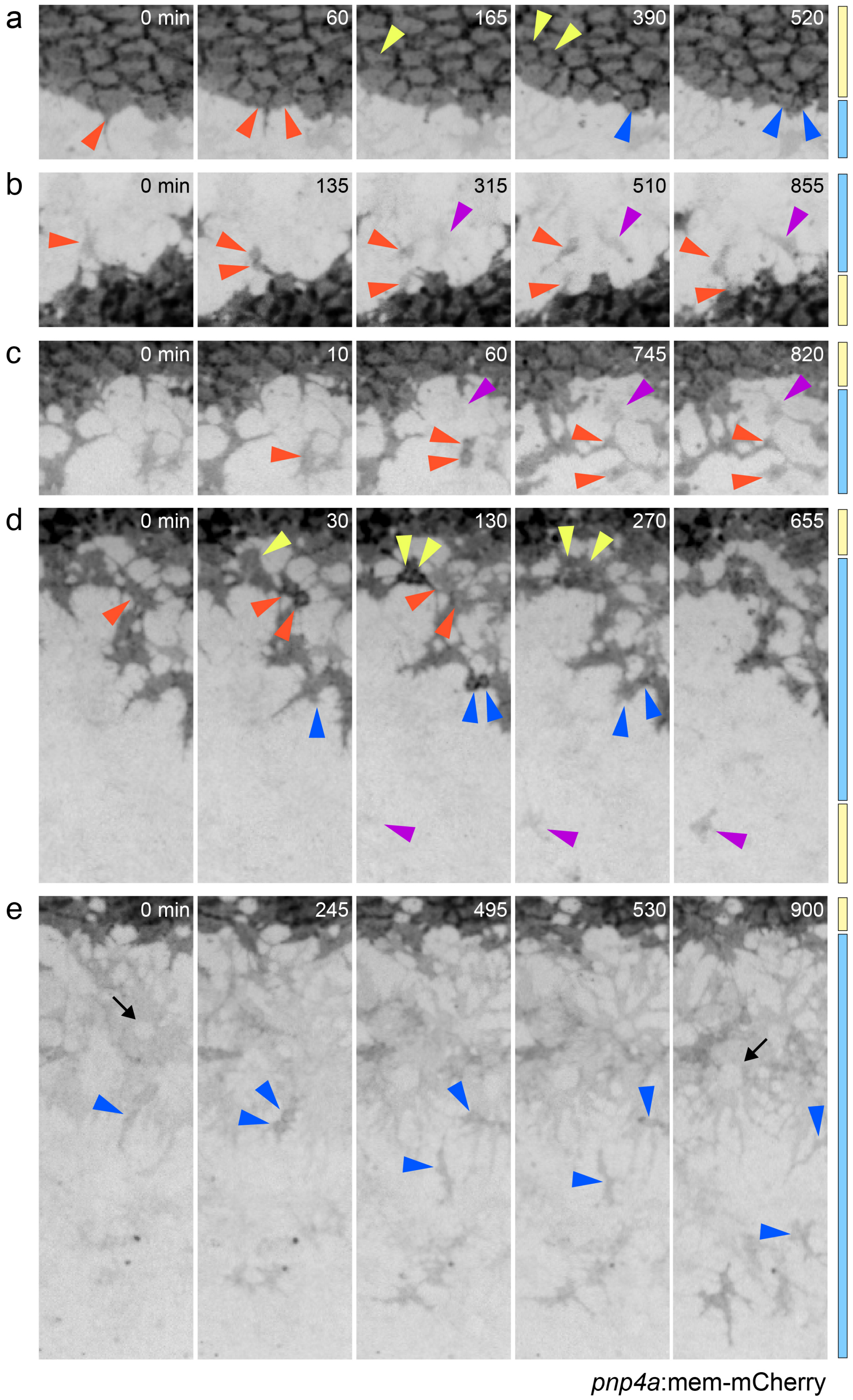

**Extended Data Figure 1. Division and migration of *pnp4a*+ cells in time-lapse imaging. (a)** Loose iridophores of stripes were more likely to divide than dense iridophores of interstripes (means±SE, *N*=27 larvae, 12,314 *pnp4*+ cells; $\chi^2$=705, *P*<0.0001, d.f.=2). **(b)** Proportions of cells having planes of division ranging from anteroposterior (0°) to dorsovental (90°) for populations within the interstripe and bounded only by other cells of the interstripe (non-edge), in the interstripe but not completely bounded by other cells of the interstripe (edge), or within the stripe. **(c)** Distances migrated (with median ± interquartile range) by loose *pnp4*+ cells of the prospective ventral primary stripe. Only cells moving greater than one half the diameter of a typical loose iridophore (~31 µm) were included. **(d)** Directions and distances migrated by same cells illustrated in C. A majority of cells moved ventrally and posteriorly from their origin. Blue circle indicates average diameter of loose iridophores.

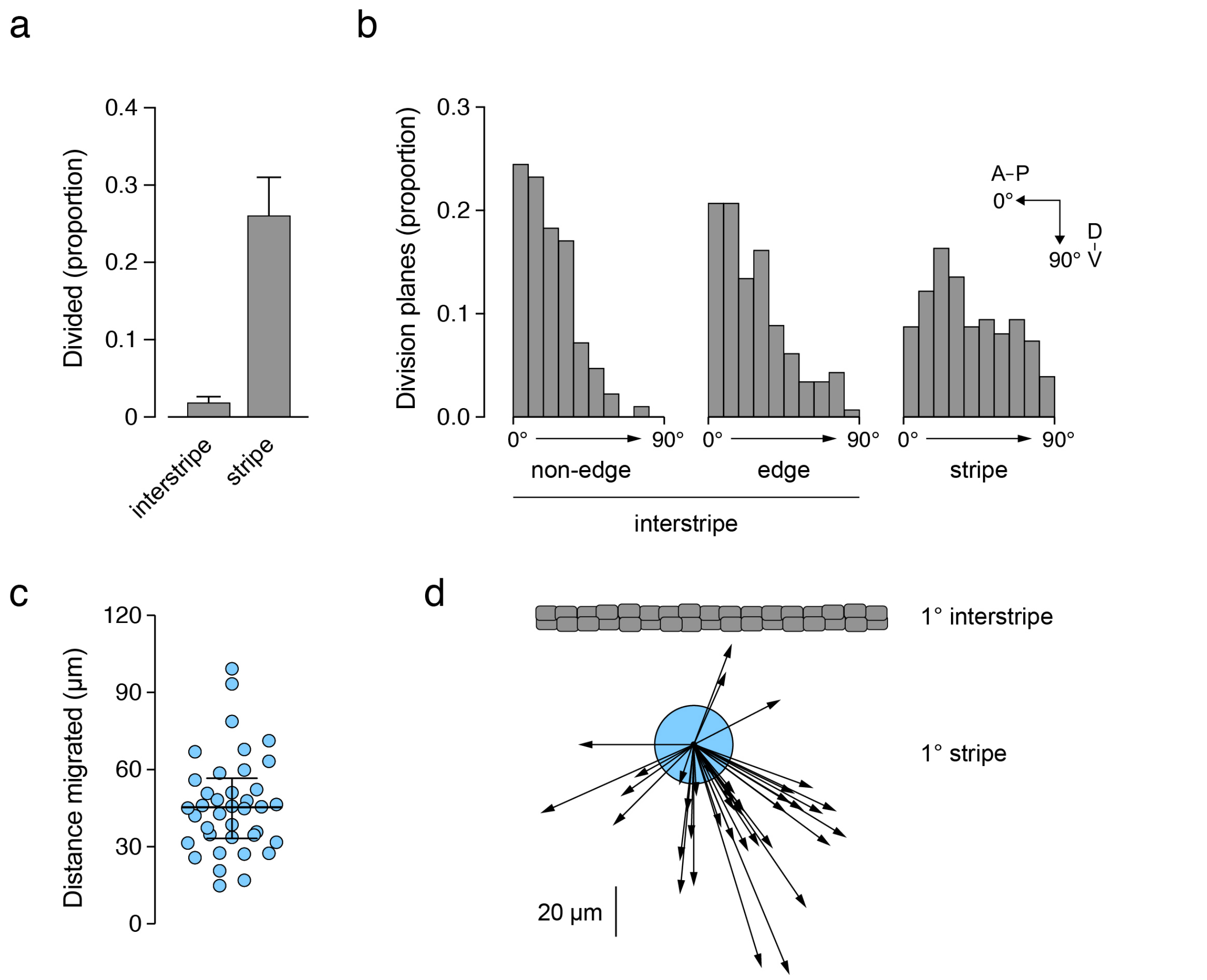


**Extended Data Figure 2. Live imaging reveals proliferation, differentiation and migration but not dynamic changes in state. (a)** *pnp4a*:mem-mCherry (inverted pixel values) reveals dividing cells at interstripe edge (orange and blue arrowheads) and within interstripe. Bars at right indicate approximate boundary between interstripe (yellow) and stripe (blue) regions and numbers in frames indicate sequential day of imaging. **(b)** Division of a cell newly expressing *pnp4a* within the prospective stripe (orange) with onset of *pnp4a* expression by an adacent cell (purple). **(c)** Division (orange) and appearance (purple) of *pnp4a*+ cells within the developing ventral primary stripe. **(d)** Division of *pnp4*+ cells (orange, yellow, blue) within stripe and appearance of new *pnp4a*+ cell (purple) at site likely corresponding to prospective secondary ventral interstripe. **(e)** Division and migration of *pnp4*+ cells within ventral stripe (blue, purple). Black arrows indicate cells having morphologies and minimal motility typical of xanthophores, which also express low levels of *pnp4a* (Saunders et al., 2019). Stages of larvae shown 7.5–8.0 SSL (Parichy et al., 2009).

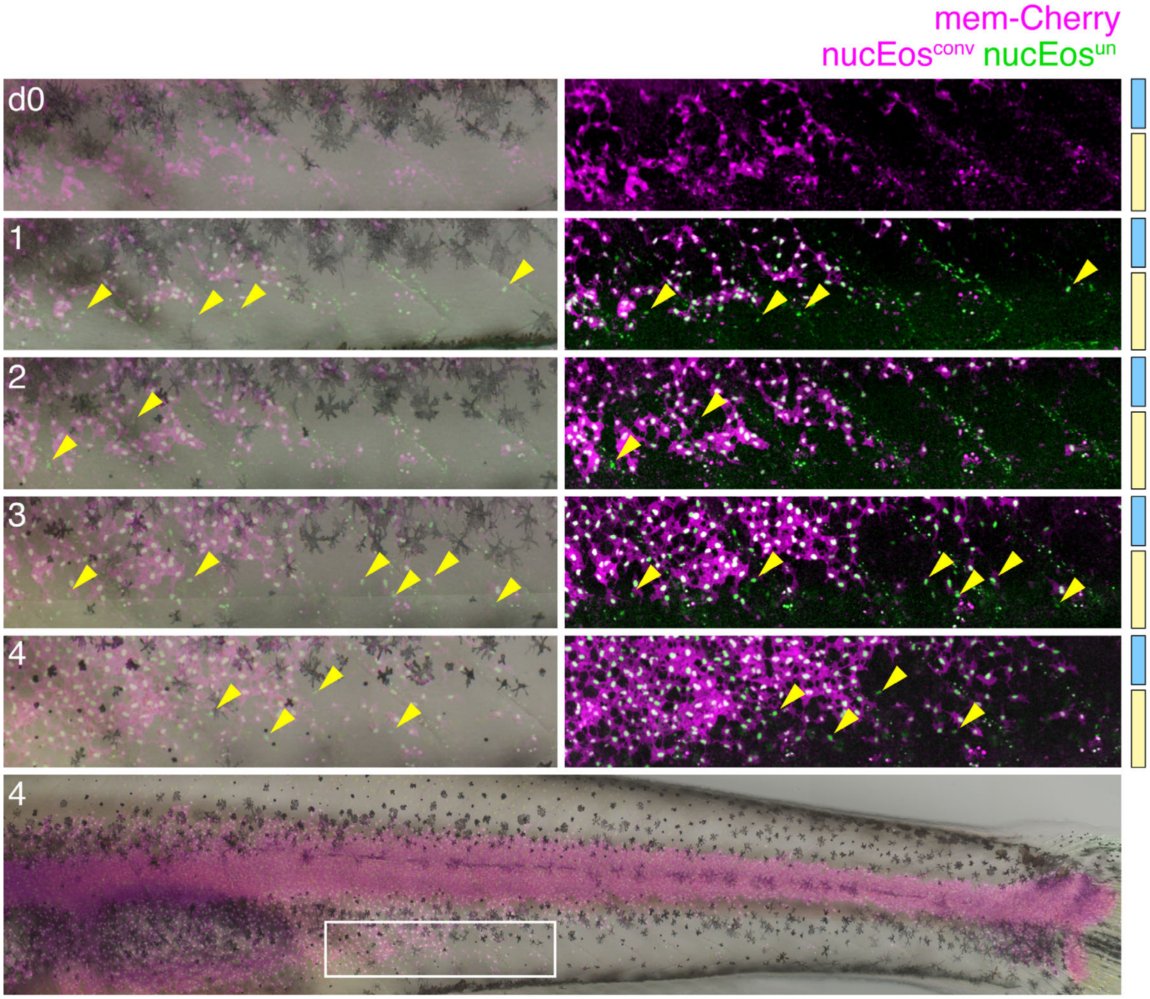


**Extended Data Figure 3. Differentiation of iridophores contibuting to secondary interstripe.** To identify newly appearing cells, whole fish were exposed to UV light daily to photoconvert all *pnp4*:nucEos from green (nucEos$^{un}$) to red (nucEos$^{conv}$; here displayed in magenta). On the following day, cells exhibiting nucEos$^{un}$ but not nucEos$^{conv}$ have arisen. Older cells typically exhibit both nucEos$^{un}$ and nucEos$^{conv}$ (white) and begin to exhibit the weaker fluorophore *pnp4*:mem-Cherry. At d0, immediately after photoconversion, all *pnp4*+ nuclei are magenta. At d1, green nuclei exhibiting only nucEos$^{un}$ are evident (e.g., arrowheads). After subsequent rounds of photoconversion, nuclei newly expressing nucEos$^{un}$ continued to appear (d2–4). Bottom panel at low magnification illustrates region highlighted above.

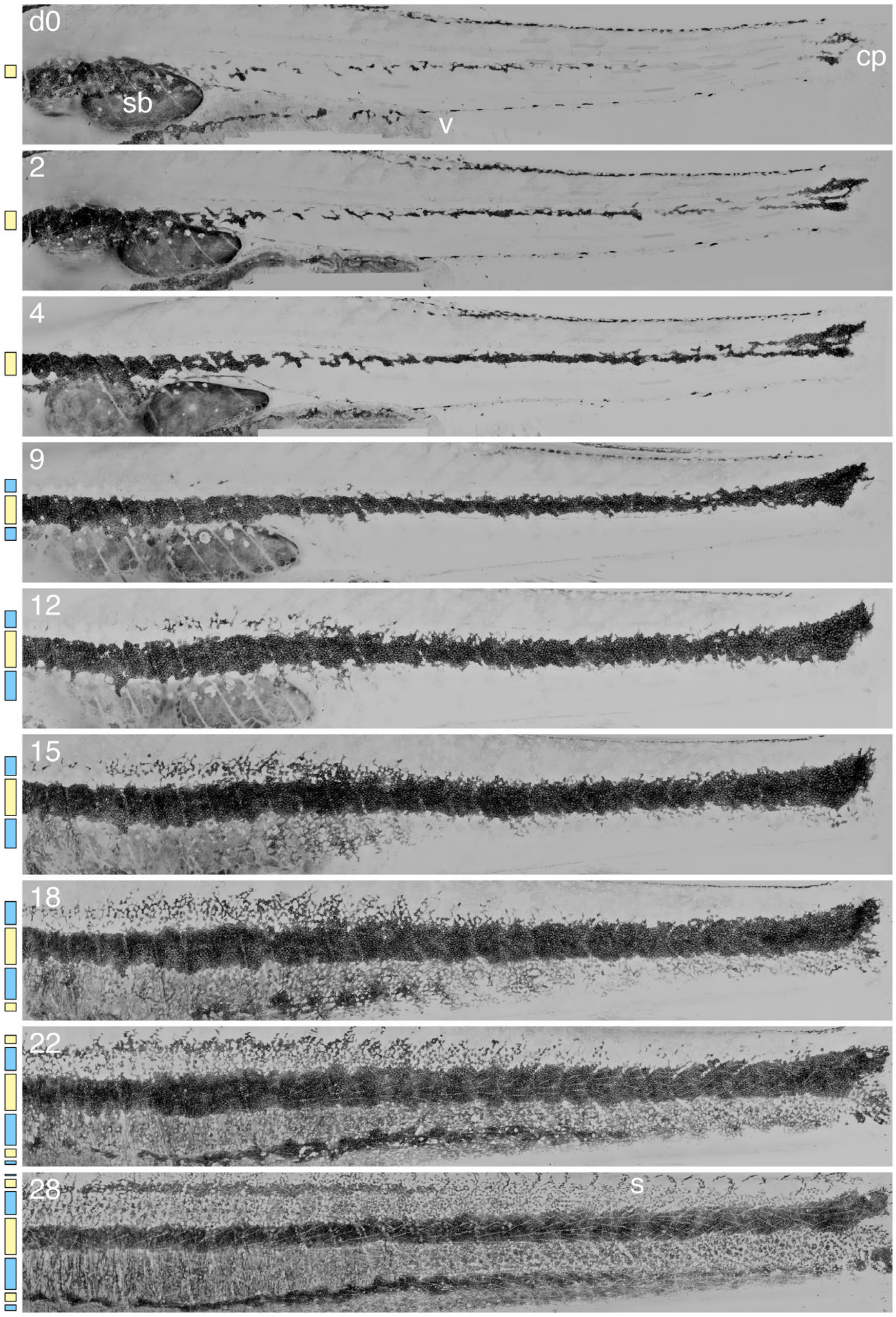


*pnp4a*:mem-mCherry

**Extended Data Figure 4. Iridophore pattern development across anteroposterior axial levels.** Shown is a representative individual imaged daily for 28 d beginning at 6.0 SSL (Parichy et al., 2009). Bars at left represent progressive appearance of *pnp4*:mem-mCherry+ cells contributing to interstripes (yellow) and stripes (blue). sb, posterior lobe of swimbladder; v, vent; cp, caudal peduncle; s, scales (d28). Images are rescaled for growth.

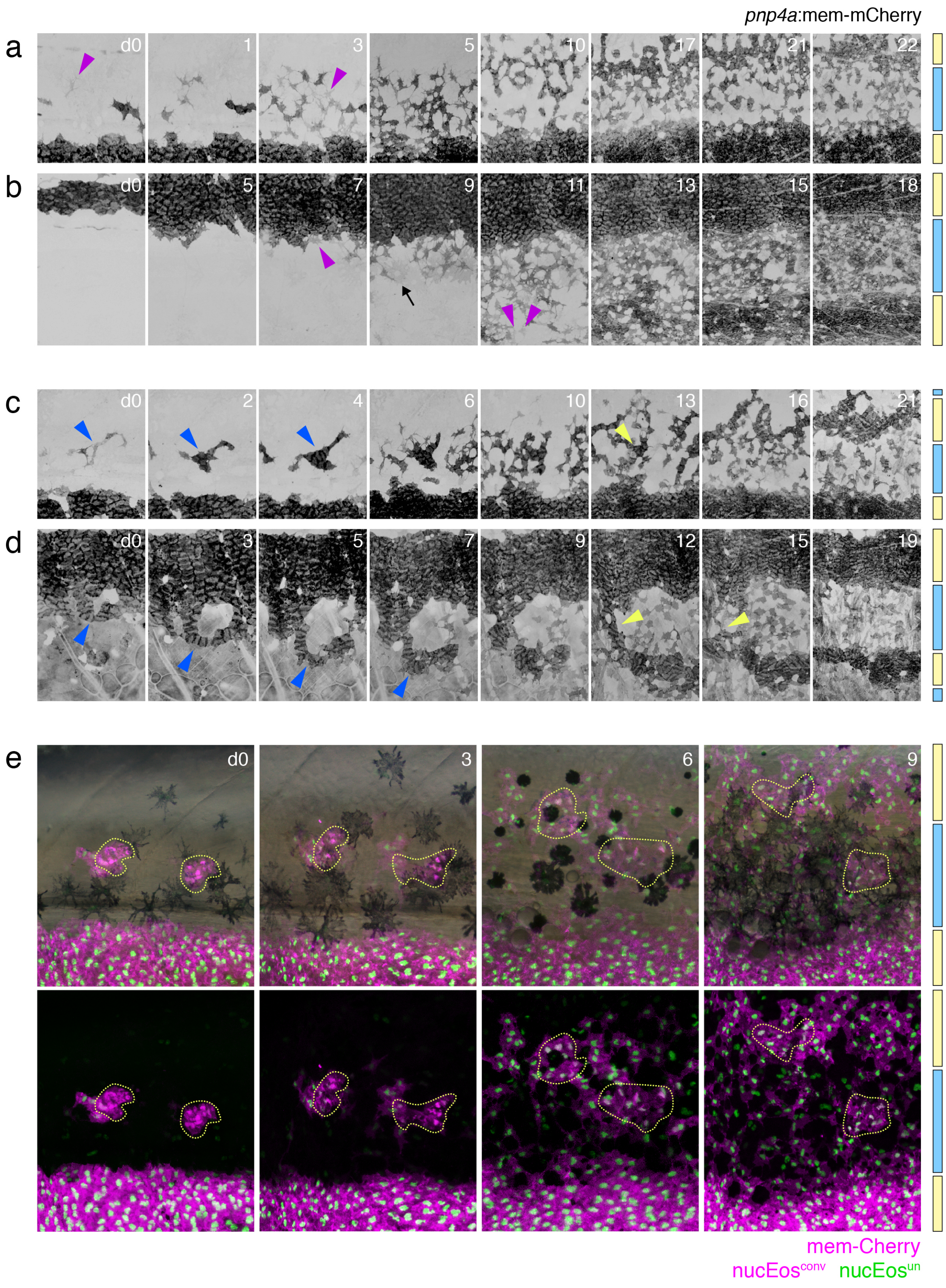

pnp4a:mem-mCherry
a
d0
1
3
5
10
17
21
22
b
d0
5
7
9
11
13
15
18
c
d0
2
4
6
10
13
16
21
d
d0
3
5
7
9
12
15
19
e
d0
3
6
9
mem-Cherry
nucEosconv
nucEosun

**Extended Data Figure 5. Repeated imaging of iridophore pattern formation and nucEos fate mapping of iridophore clusters within the dorsal stripe. (a–d)** Details from stitched images that covered the anterior–posterior and dorsal–ventral boundaries of pattern formation on the body, examples of which are provided in Supplementary Movies S6 and S7. Bars at right indicate prospective interstripe (yellow) and stripe (blue) regions. **(a,b)** Appearance of *pnp4a*+ cells (purple arrowheads) within the dorsal stripe (a) and ventral stripe (b), and emergence of the secondary interstripes. Black arrow indicates cell with typical xanthophore morphology, weakly expressing *pnp4a*:mem-mCherry. **(c,d)** Examples of patterning observed in a minority of wild-type fish. In (c), several iridophores arise in an initially dense arrangmenet (blue arrowheads) within the prospective dorsal stripe and subsequently assumes a more dispersed arrangement (yellow arrowhead) within the stripe. In (d), a ventrally extending group of densely packed iridophores (blue arrowheads) ultimately becomes separated from the primary interstripe (yellow arrowheads) and contributes to the second ventral interstripe. **(e)** Photoconversion of *pnp4a*:nucEos expressed by iridophores in small clusters within the developing dorsal stripe revealed cells retaining photoconverted fluorophore (white nuclei) within the dorsal interstripe (left cluster) and within the dorsal stripe (right cluster). Some additional cells within the marked regions began expressing *pnp4a*:nucEos only after photoconversion (green nuclei).

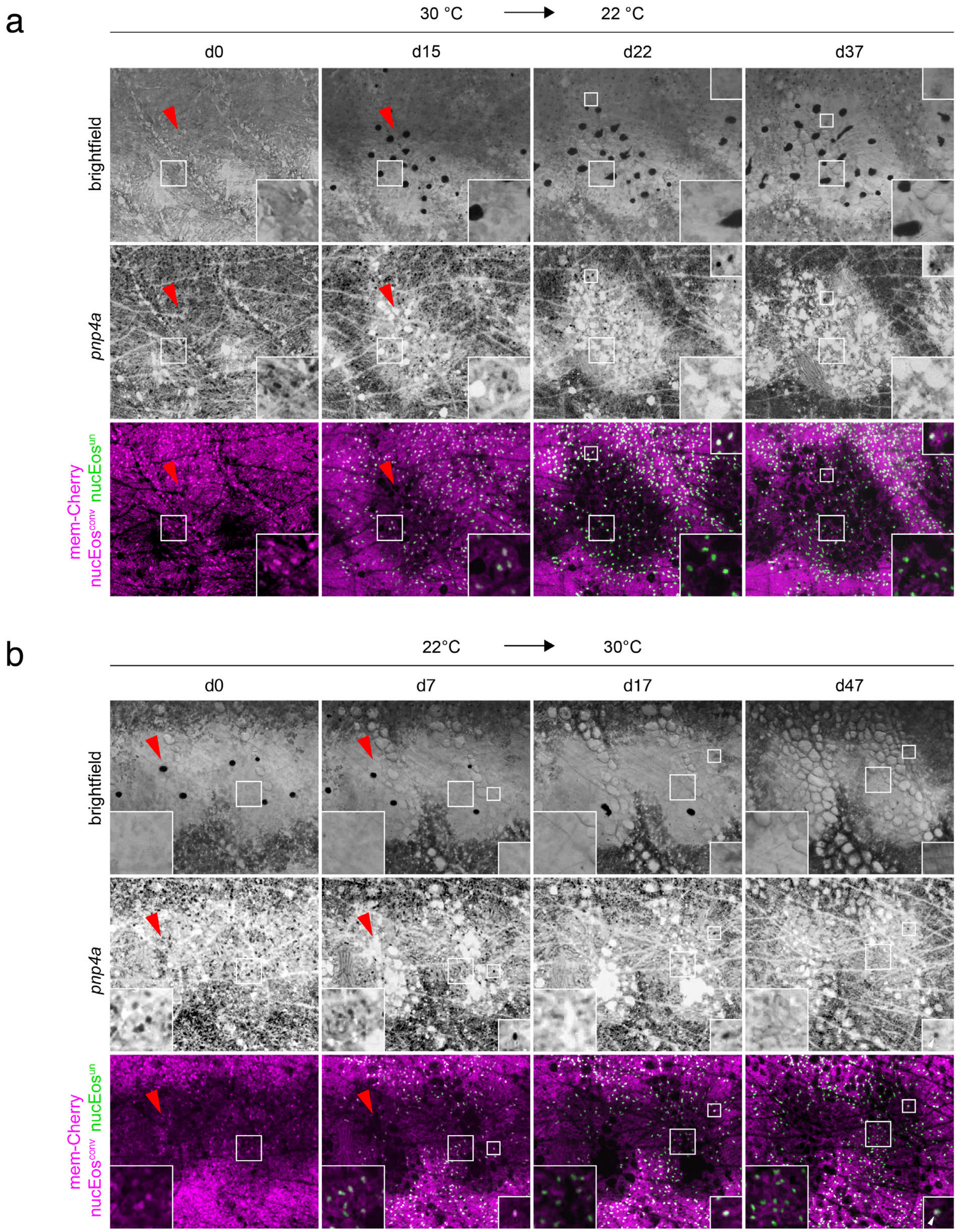
a
30 °C
22 °C
d0
d15
d22
d37
brightfield
pnp4a
mem-Cherry
nucEos^conv nucEos^un
b
22°C
30°C
d0
d7
d17
d47
brightfield
pnp4a
mem-Cherry
nucEos^conv nucEos^un

**Extended Data Figure 6. Differentiation of new iridophores during pattern remodeling.** Shown are details of repeatedly imaged, representative adult fish homozygous for the temperature sensitive allele *mitfa*$^{vc7}$ . Fish were stably transgenic to express *pnp4a*:mem-mCherry in all iridophores, marking cell outlines, and were transiently transgenic for *pnp4a*:nucEos, thereby expressing photoconvertible fluorophore mosaically among iridophores. nucEos was photoconverted and fish were then shifted between temperatures to promote melanophore development or death. **(a)** Shift from restrictive temperature (30°C) to permissive temperatures (22 °C) allowed melanophore differentiation by ~d2, continuing through ~d22 (upper panels, brightfield). The appearance of melanophores was followed by a gradual remodeling of iridophore pattern from a relatively dense to a looser arrangement (middle panels, *pnp4a*). Photoconverted nuclei (magenta; e.g., lower panel, inset on d0) gradually acquired unconverted nucEos (white; inset on d15), but many were subsequently lost, accompanied by the appearance of spaces devoid of iridophores (arrowheads). Later, cells expressing only unconverted nucEos increased in abundance (green; e.g., large insets on d22, d47) though other cells retaining photoconverted nucEos remained evident at margins of remodeled regions (small insets on d22, d37). **(b)** Reciprocal temperature shift caused the death of melanophores (Lewis et al., 2019) and iridophore pattern remodeling from a loose to denser arrangement. Remodeling proceeded through a period in which regions became devoid of iridophores with photoconverted nucEos (arrowheads). Although some iridophores with photoconverted nucEos persisted through d47 (white; small insets d7–47, and some cells in large inset on d7), iridophores populating regions without melanophores were increasingly likely to exhibit only unconverted nucEos (green; large insets d17, d47), suggesting they were newly differentiating.

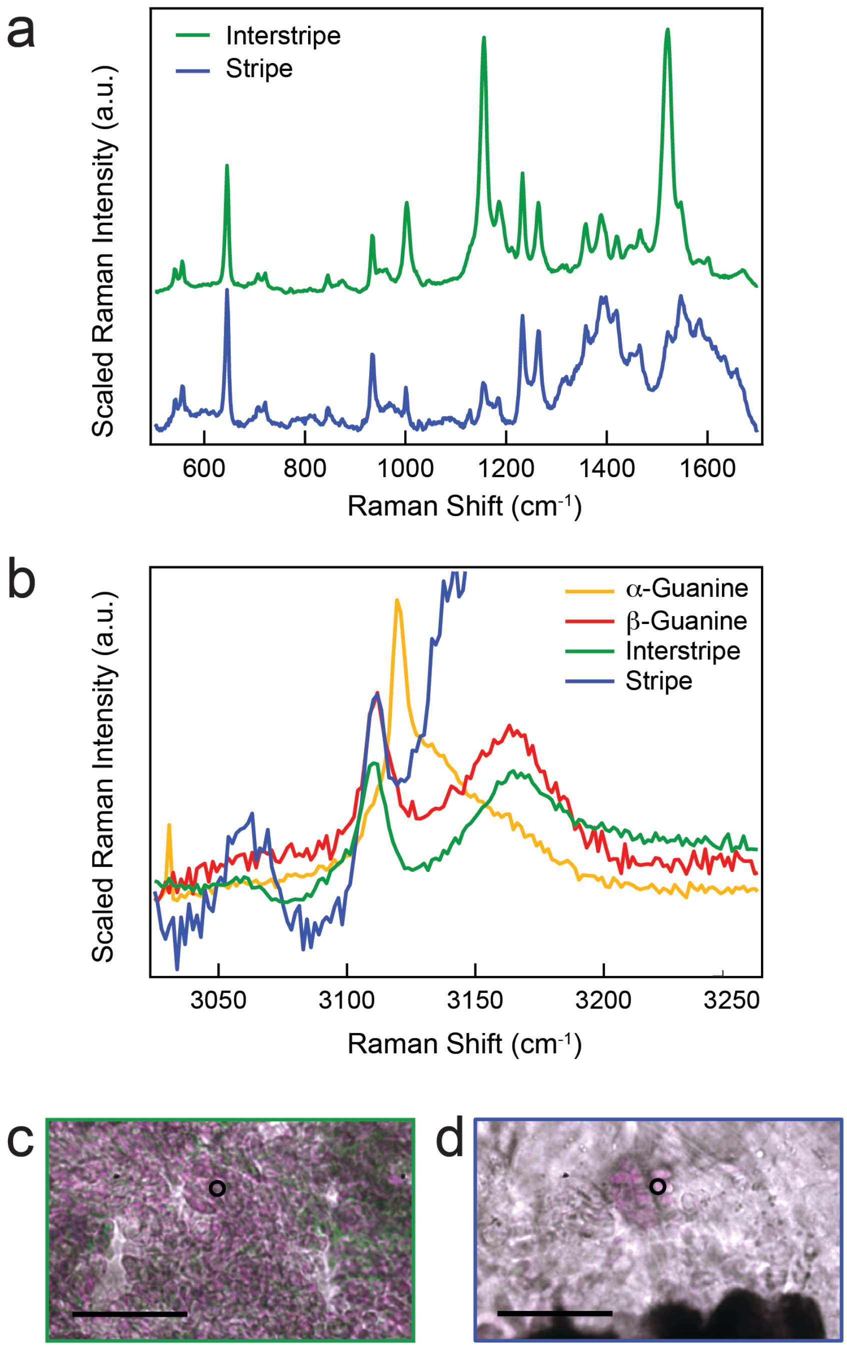


**Extended Data Figure 7. Raman micro-spectroscopy of zebrafish stripe and interstripe iridophores. (a)** Representative Raman spectra taken from interstripe (green; n=9 cells) and stripe (blue; *n*=14) cells in zebrafish tissue. **(b)** Brightfield images of interstripe (green) and stripe (blue) cells. Black circles indicate collection location of Raman spectra in (a). Scale bar = 10 μm. **(c)** High energy peak pattern (dashed line) indicates β-guanine is present in both interstripe and stripe cells.

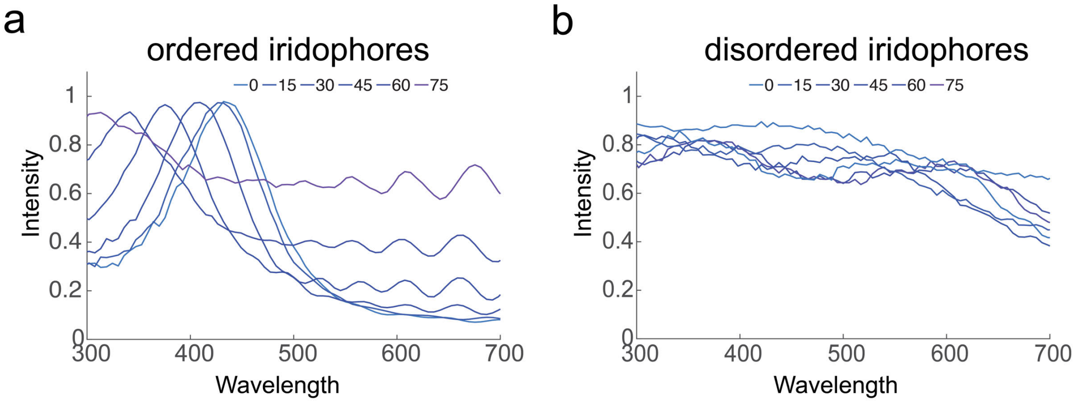


**Extended Data Figure 8. Simulations of the expected reflection from iridophores at different angles of incident light. (a,b)** Monte Carlo based simulations of the reflection expected from ordered (a) and disordered iridophores (b) at different angles of incident light (0-75°). While there was very little effect over the reflection of the disordered iridophores (b), a clear angular dependence was observed for the ordered iridophores (a) showing a blue shift in the reflected light with increasing angle of incident light.

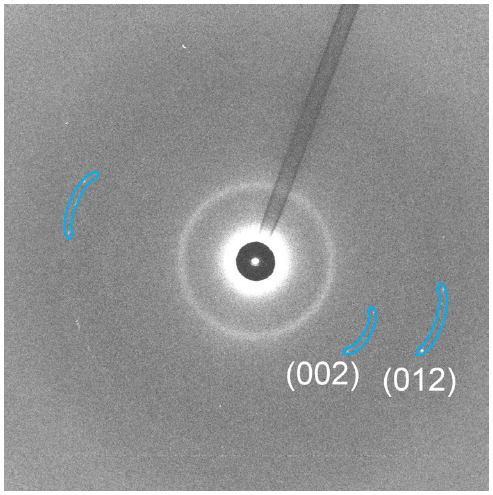


**Extended Data Figure 9. Micro X-ray diffraction of an ordered iridophore from the posterior flank of a *mitfa*$^{w2}$ (nacre) fish.** In this X-ray diffraction, low angular distribution diffractions of both the (012) and the (002) planes, typical of the ordered iridophores were observed.

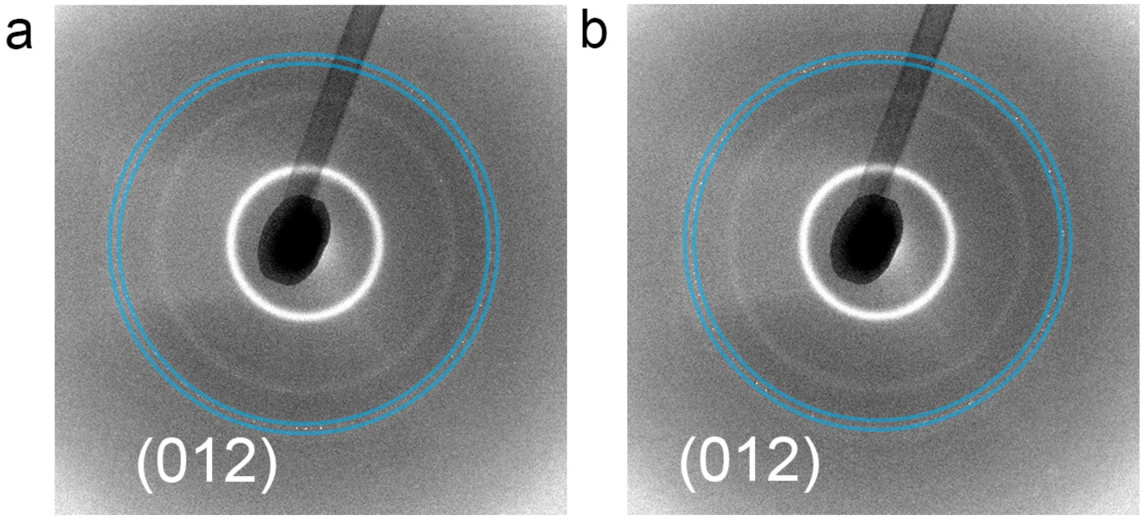


**Extended Data Figure 10. Micro X-ray diffraction of a 6.5 SSL larva. (a,b)** X-ray diffraction patterns from a vertical line measured across the trunk of the fish, only high angular distributions of the (012) plane were observed.

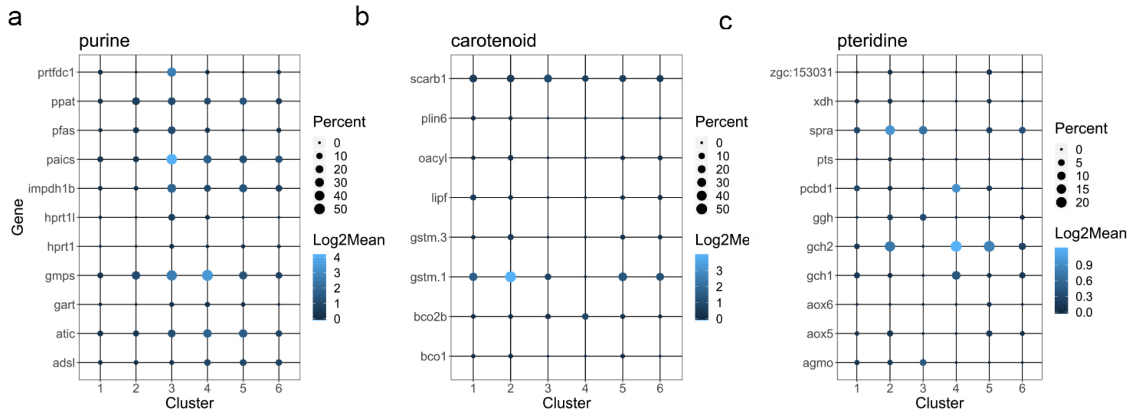


**Extended Data Figure 11. Purine metabolism pathways. (a-c)** scRNA-Seq analysis showing aggregate expression profiles per cluster for genes associated with different metabolism pathways; (a) purine (b) carotenoid (c) pteridines.

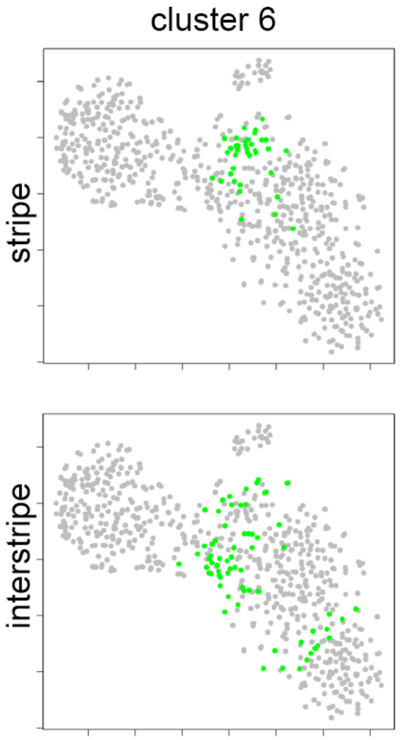


**Extended Data Figure 12.** Two-dimensional tSNE representation of the collected skin cell clusters showing the anatomical origin of cluster 6 iridophores is split between the stripe (33%) and the interstipre (67%).

**Supplementary Movies legends-**

**Supplementary Movie S1. Proliferation of densely arranged *pnp4*+ cells.** Region shown corresponds to that described in Figure 2A with arrowheads illustrating cell division both at the edge of the interstripe and within it.

**Supplementary Movie S2. Proliferation and differentiation of *pnp4*+ cells adjacent to interstripe.** Region shown corresponds to that of Figure 2B, with orange arrowheads indicating proliferative cell and daughters.

**Supplementary Movie S3. Proliferation and differentiation of loosely arranged *pnp4*+ cells within the prospective stripe.** Region shown includes that in Figure 2C. Orange arrowheads indicate proliferative cell and daughters, purple arrowhead indicates cell newly expressing *pnp4a*:mem-mCherry.

**Supplementary Movie S4. Proliferation and differentiation of loosely arranged *pnp4*+ cells within the prospective stripe.** Region shown corresponds to that of Figure 2D. Arrowheads indicate proliferating cells and daughters except for purplke arrowhead that marks a ventral cell newly acquiring *pnp4a*:mem-mCherry expression.

**Supplementary Movie S5. Proliferation and migration of *pnp4*+ cells within stripe.** Region corrresponds to Figure 2E, with arrowheads identifying a cell that divides into daughter cells that become widely separated after one migrates ventrally.

**Supplementary Movie S6. Iridophore patterning from anterior to posterior.** Movie shows a typical individual, imaged daily from stages of primary interstripe development through stripe formation and interstripe reiteration. Individual corresponds to that shown in Supplementary Figure S4A,B. Images are rescaled and aligned to control for growth.

**Supplementary Movie S7. Iridophore patterning from anterior to posterior.** Movie shows an individual that represents a minority of instances, with transiently dense clusters of iridophores in the prospective dorsal stripe, as well as a patche of densely arranged iridophores that splits ventrally between primary and second interstripe. Movie corresponds to individual shown in Supplementary Figure S4C,D. Images are rescaled and aligned to control for growth.

**Supplementary Movie S8. The response of the stripes patent to NE.** Movie shows the stripe pattern response of an adult zebrafish to NE. Minutes after administrating NE the contrast between the dark blue stripe and the yellow interstripe is drastically diminished as the melanophores aggerated and the stripe iridophores changed their color from blue to yellow.
